# The resource elasticity of control

**DOI:** 10.1101/2024.12.16.628674

**Authors:** Levi Solomyak, Aviv Emanuel, Eran Eldar

## Abstract

The ability to determine how much the environment can be controlled through our actions has long been viewed as fundamental to adaptive behavior. While traditional accounts treat controllability as a fixed property of the environment, we argue that real-world controllability often depends on the effort, time and money we are able and willing to invest. In such cases, controllability can be said to be *elastic* to invested resources. Here we propose that inferring this elasticity is essential for efficient resource allocation, and thus, elasticity misestimations result in maladaptive behavior. To test this hypothesis, we developed a novel treasure hunt game where participants encountered environments with varying degrees of controllability and elasticity.

Across two pre-registered studies (N=514), we first demonstrate that people infer elasticity and adapt their resource allocation accordingly. We then present a computational model that explains how people make this inference, and identify individual elasticity biases that lead to suboptimal resource allocation. Finally, we show that overestimation of elasticity is associated with elevated psychopathology involving an impaired sense of control. These findings establish the elasticity of control as a distinct cognitive construct guiding adaptive behavior, and a computational marker for control-related maladaptive behavior.

## Introduction

Taking action requires us to invest resources, such as time, money, and effort, and is thus only sensible if our actions have the potential to change the environment to our advantage. Accurate estimation of this potential – that is, of the environment’s controllability^1–4^ – enables us to make informed decisions as to whether to engage in action^5–8^. Misestimations of controllability thus result in maladaptive behavior, and contribute to mental health disorders such as depression, anxiety, and obsessive-compulsive disorder^9–13^.

The degree of control we possess over our environment, however, may itself depend on the resources we are willing and able to invest. For example, the control a biker has over their commute time depends on the power they are willing and able to invest in pedaling. In this respect, a highly trained biker would typically have more control than a novice. Likewise, the control a diner in a restaurant has over their meal may depend on how much money they have to spend. In such situations, controllability is not fixed but rather *elastic* to available resources (i.e., in the same sense that supply and demand may be elastic to changing prices^14^). To formalize how elasticity relates to control, we build on an established definition of controllability as the fraction of reward that is controllably achievable^15^, χ. Uncertainty about this fraction could result from uncertainty about the amount of resources that the agent is able and willing to invest, max C. Elasticity can thus be defined as the amount of information obtained about controllability by knowing the amount of available resources: I(χ; max C).

While only controllable environments can be elastic, the inverse is not necessarily true – controllability can be high, yet inelastic to invested resources – for example, choosing between bus routes affords equal control over commute time to anyone who can afford the basic fare (Figure 1; Supplementary Note 1). That said, since all actions require some resource investment, no controllable environment is completely inelastic when considering the full spectrum of possible agents, including those with insufficient resources to act (e.g., those unable to purchase a bus fare or pay for a fixed-price meal). Even in this case, however, controllability can be elastic to varying degrees.

**Figure 1.**
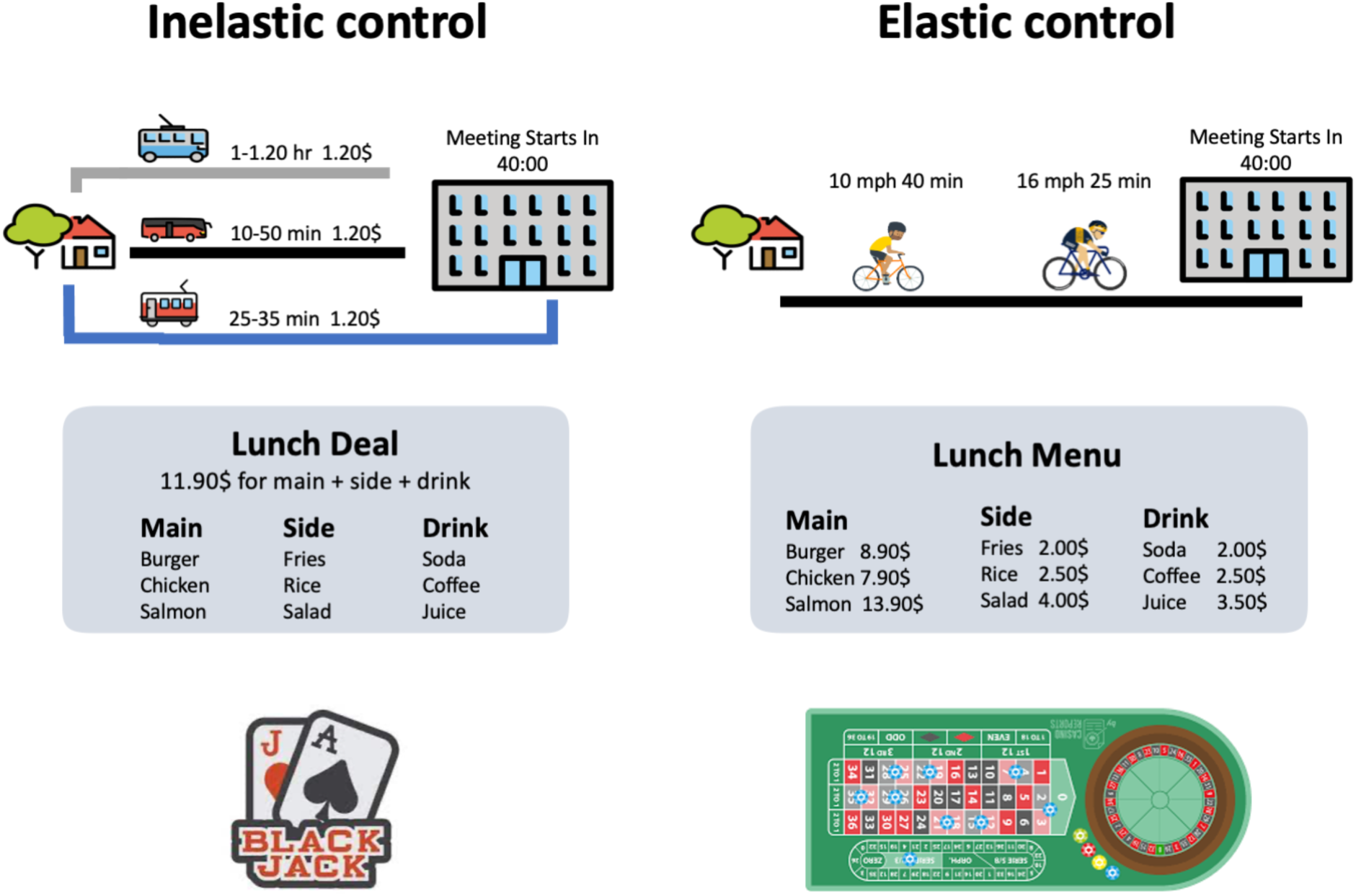
Examples of inelastic vs elastic control: Top –. Choosing one of three equally priced public transport routes provides inelastic control over commute time, as control does not increase with additional available resources beyond the basic fare. In contrast, the control of a biker over commute time is elastic to the power they can invest in pedaling. **Middle –** A diner has either inelastic or elastic control over their lunch depending on whether the restaurant offers a fixed-price lunch deal where all diners have the same options, or a standard menu where more expensive dishes are only available to those able to spend more. **Bottom –** the probability of winning in blackjack is inelastic to money, as it only depends on whether one hits or stands, whereas when playing roulette, one can increase the probability of winning by investing more coins to cover more possible outcomes. Importantly, most real-world scenarios lie on a spectrum between these illustrative extremes, such that controllability is partly elastic and partly inelastic. In all situations, agents must infer the degree to which controllability is elastic to be able to determine whether the potential gains in control outweigh the costs of investing additional resources (e.g., physical exertion, money spent, time invested).

Knowing the elasticity of control is necessary for deciding whether and how to act. Failing to achieve one’s goal, for example, should only lead one to invest more resources in an elastic environment. Critically, in many real-life domains (e.g., studying for an exam), we must use our experiences to infer the degree to which we can improve our control by investing more resources^16–22^. Failure to do so is bound to result in maladaptive behavior, and may thus contribute to mental health disorders. Depressed patients, for instance, are less willing to expend their resources to obtain control^23,28–29^, which could reflect an underestimation of elasticity. Conversely, the repetitive and unusual amount of effort invested by people with obsessive-compulsive disorder in attempts to exert control^25,26^ could indicate an overestimation of elasticity, that is, a belief that adequate control can only be achieved through excessive and repeated resource investment^26^. Ultimately, not having a good grasp of how additional efforts influence the success of one’s actions is bound to undermine one’s sense of agency^16–17^.

Thus, here we ask whether and how people infer the elasticity of control over their environment. We hypothesize that individual differences in elasticity inference lead to a mismanagement of one’s resources when attempting to control the environment, and are thus associated with psychopathologies involving a distorted sense of control^23–26^.

Experimental paradigms to date have conflated overall controllability and its elasticity, such that controllability was either low or elastic^16–22^. The elasticity of control, however, must be dissociated from overall controllability to accurately diagnose mismanagement of resources. A given individual, for instance, may tend to overinvest resources because they overestimate controllability – for example, exercising due to a misguided belief that this can make one taller, when in fact height cannot be controlled. Alternatively, they may do so because they overestimate the elasticity of control – for example, a chess expert practicing unnecessarily hard to win against a novice, when their existing skill level already ensures control over the match’s outcome. Identifying which misestimation drives resource mismanagement requires an experimental design that independently manipulates overall controllability from its elasticity.

Therefore, here we dissociate elastic and inelastic controllability using a novel treasure-hunt game where participants encountered three distinct environments: The first offered *high elastic controllability*, where a high level of control can only be attained by investing extra resources. The second offered *high inelastic controllability*, where control is attainable with any level of resource investment. And the third offered *low controllability*, where control was not attainable despite any level of resource investment. In each environment, participants were allowed to invest various amounts of resources to pursue reward or to altogether forgo pursuing reward. Examining how participants adapted their resource investment to observed outcomes enabled us to determine whether and how participants inferred each environment’s controllability and its elasticity.

Across two pre-registered studies (N = 514), we establish that people do indeed infer the elasticity of control. We present a computational model that explains how people make this inference, and identify individual biases in its implementation. We thus show that elasticity overestimation leads to an overinvestment of resources and is associated with elevated psychopathology involving an impaired sense of control. These findings establish elasticity as a crucial dimension of controllability that guides adaptive behavior, and a computational marker of control-related psychopathology.

## Results

We had online participants (first study = 264, replication study = 250, ages =18-45, Mean 33 ± .6) play a novel treasure-hunt game set across multiple planets. On each trip (trial), participants attempted to travel from an initial location (‘desert’ or ‘fountain’) to a treasure (worth 150 coins; 15¢ of monetary compensation), located at either the house or the mountain (Figure 2A). Participants could exercise control by boarding the train to reach the house, or the plane to reach the mountain (right-side Figure 2B). The present planet’s controllability determined the probability that participants would succeed in boarding the vehicle they selected. In high-controllability planets, participants consistently boarded their preferred vehicle, enabling them to reliably reach their destination. Conversely, in low-controllability planets, participants frequently failed to board and were thus forced to walk to the nearest location. On 20% of trials, the nearest location was where the treasure was located, thus partially decoupling success in exerting control from reward.

**Figure 2.**
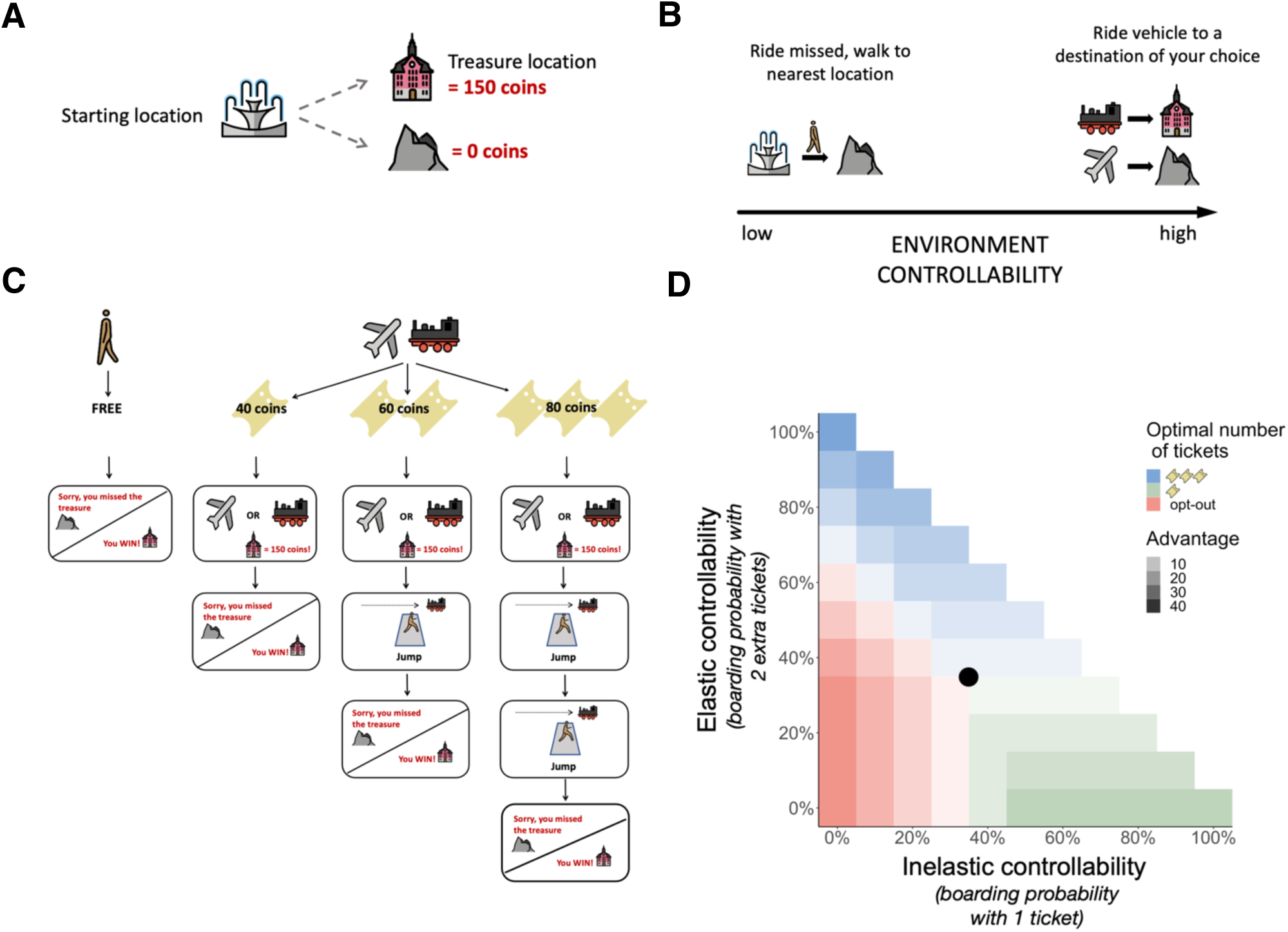
Experimental design: (A) **Goal.** On each trip to a planet, participants’ goal was to reach a treasure located either at the house or the mountain (B) **Transition rules.** Participants could exercise control by boarding either the plane or the train to a destination of their choice, whereas missing the ride sent the participant walking to the nearest location (which happened to be the treasure location in only 20% of trials). (C) **Trip structure.** At the beginning of each trip, participants selected whether to purchase 1, 2, or 3 tickets to attempt to board their vehicle of choice, or walk for free to the nearest location. If the participant purchased at least one ticket, they were allowed to choose between the plane and the train. Then, for each additional ticket purchased, the participant was given an opportunity to increase their chances of boarding the vehicle. Specifically, the chosen vehicle appeared moving from left to right across the screen, and the participant attempted to board it by pressing the spacebar when it reached the center of the screen. At the end of the trip, participants were shown where they arrived at, allowing them to infer whether they successfully boarded the vehicle. (D) **The space of possible planets.** Planets varied in inelastic controllability – the probability that even one ticket would lead to successfully boarding the vehicle – and in elastic controllability – the degree to which two extra tickets would improve the probability of successfully boarding the vehicle (one extra ticket provided half the benefit). The color corresponds to the optimal number of tickets (0, 1 or 3) in each planet. The darker the color the higher the advantage in expected value gained by purchasing the optimal number of tickets relative to the second-best option. Each participant made 30 consecutive trips in one planet from the green area, one planet from the blue area, one planet from the red area, and one planet with identical characteristic across all participants (black circle).

To investigate how people learn the elasticity of control, we allowed participants to invest different amounts of resources in attempting to board their preferred vehicle. Participants could purchase one (40 coins), two (60 coins), or three tickets (80 coins) or otherwise walk for free to the nearest location. Participants were informed that a single ticket allowed them to board only if the vehicle stopped at the station, while additional tickets provided extra chances to board even after the vehicle had left the platform. For each additional ticket, the chosen vehicle appeared moving from left to right across the screen, and participants could attempt to board it by pressing the spacebar when it reached the center of the screen. Thus, each additional ticket could increase the chance of boarding but also required a greater investment of resources—decreasing earnings, extending the trial duration, and demanding attentional effort to precisely time a button press when attempting to board.

Participants received no feedback about individual boarding attempts but were shown their initial and final locations at the end of each trip, enabling them to infer whether they had successfully boarded their vehicle. For simplicity, we defined inelastic controllability as the probability that even one ticket would lead to successfully boarding the vehicle, and elastic controllability as the degree to which two extra tickets would increase that probability.

To dissociate elastic from inelastic controllability, each participant visited three randomly sampled planets. In the first planet, controllability was high and inelastic, meaning that no matter how many tickets you purchased, you had a good chance of making your ride (sampled from green area in Figure 2D). In the second planet, controllability was high and elastic, meaning that you needed to purchase two extra tickets to have a good chance of making your ride (Figure 2D, blue). And in the third planet, controllability was low, meaning that no matter how many tickets you purchased, you did not have a good chance of making your ride (Figure 2D, red).

Importantly, the order of these planets was randomized, and participants did not know in advance which planet they were visiting. Thus, they could only infer a planet’s controllability and elasticity from the outcomes of their actions. Moreover, to increase sensitivity to individual biases, we had all participants visit a fourth planet where purchasing any number of tickets (0, 1, 2, 3) was equally optimal (Figure 2D, black circle). We also homogenized participants’ initial learning experiences by giving them three free tickets for the first five visits to every planet. As we shall see below, this latter measure also helped dissociate between different models of how participants solved the task.

### People adapt to the elasticity of control

To determine whether participants inferred the elasticity of control, we first examined how they adapted their resource investment across planets. We found that participants were more likely to purchase tickets (‘opt in’) in more controllable planets, whether controllability was elastic (Initial and Replication studies, *p*<.001; mixed logistic regression) or inelastic (Initial and Replication studies, *p* <.001; Figure 3A&C). Conversely, participants purchased more extra tickets only in planets with high elastic controllability (Initial *p* =.01, Replication *p*=.004; mixed probit regression; Figure 3B&C). Although substantial individual differences were observed in this regard (Figure 3D), these results show that most participants successfully adapted to both the controllability of the environments they visited and its elasticity.

**Figure 3.**
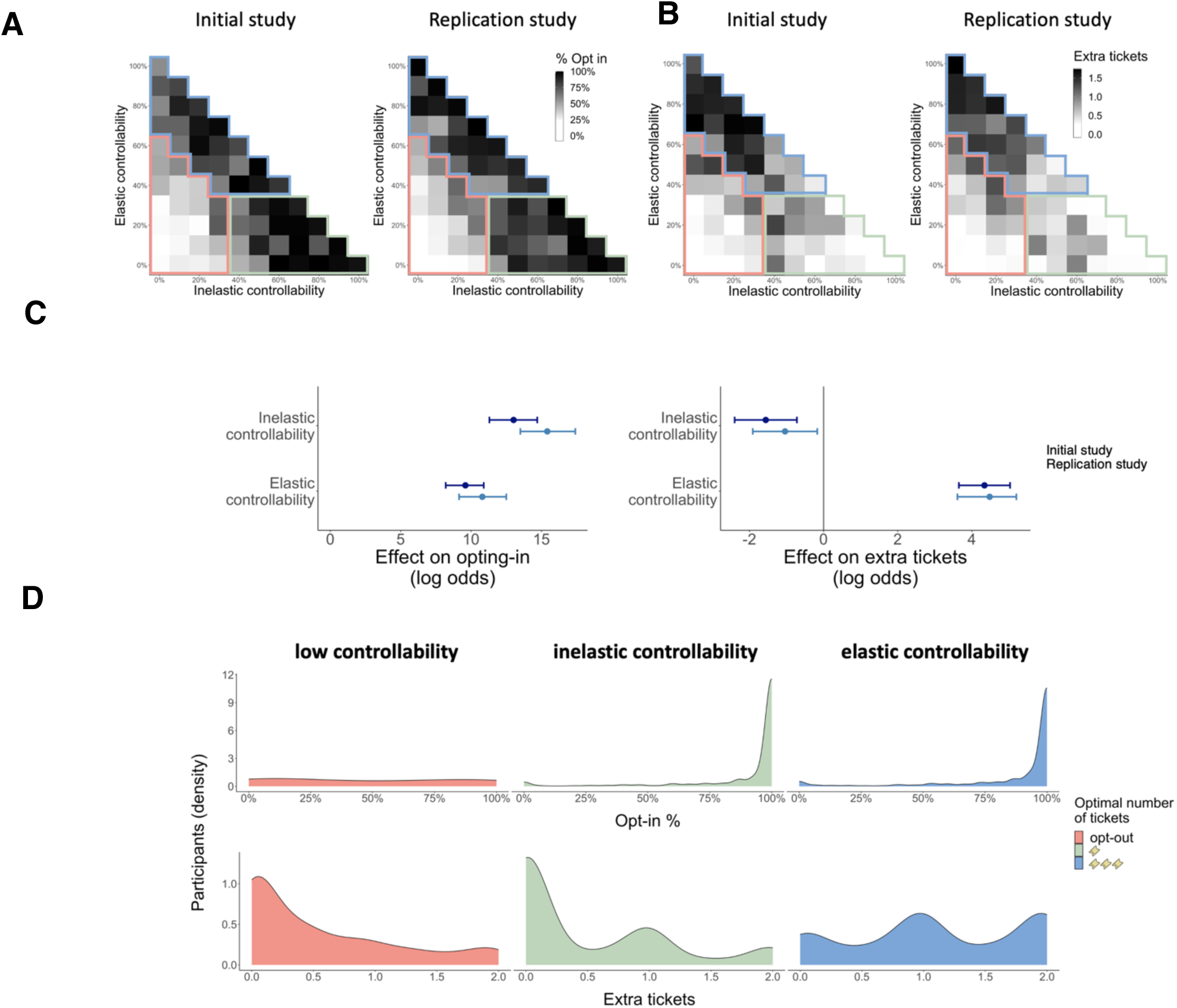
Participants adapted to the elasticity of control: Results from initial (N = 264) and replication (N = 250) studies. (A) **Opt-in percentage across all planets.** Participants opted-in more frequently on controllable planets (outlined in blue and green) than on uncontrollable planets (outlined in red) (B) **Extra tickets purchased across all planets.** Participants purchased more extra tickets in planets with high elastic controllability (outlined in blue). An average of 11 to 12 participants visited each planet in each of the studies. (C) **Effect of elastic and inelastic controllability on opting in and the purchasing of extra-tickets.** Bars show estimated fixed effects and 95% CIs from mixed logistic (opt-in) and probit (extra tickets) regressions. (D) **Individual differences.** Distribution of participants by opt-in rate (top) and average number of extra tickets purchased (bottom) across both studies, shown separately for planets with low controllability (red), high inelastic controllability (green), and high elastic controllability (blue).

### People infer the elasticity of control

That participants adapted to each environment does not necessarily mean they inferred the elasticity of control (nor that they inferred control at all, but see Methods for how this was verified). Successful adaptation to the elasticity of control may also be achieved merely by treating each possible level of resource investment as a separate course of action. One could thus learn the degree of control afforded by each course of action (i.e., purchasing 1, 2, or 3 tickets), and then choose the one that affords the highest expected value considering its cost. This simple strategy could achieve optimal performance given unlimited experience. However, it would not be as efficient as a strategy that employs elasticity inference, because it does not make use of the dependencies that exist between different levels of resource investment in the degrees of control they afford. Thus, upon failing to board a vehicle despite purchasing 3 tickets, the simpler strategy would need to purchase 1 or 2 tickets to know these options would fail as well, whereas an agent equipped with the concept of elasticity would conclude that controllability is low, and it is thus best to opt out. Conversely, upon succeeding to board with 3 tickets, the simpler strategy would tend to favor purchasing 3 tickets, over 1 or 2 tickets, even if the extra tickets confer no benefit, whereas an agent that infers elasticity would not favor 3 over 1 or 2 tickets because it received no evidence that controllability is elastic (i.e., that the extra tickets help).

To determine which of these models best describes the way participants solved the task, we formalized the simpler strategy as the ‘controllability model’, and an elasticity-inferring strategy as the ‘elastic controllability model’ (pre-registered at: https://aspredicted.org/CHW_12H). Both models use beta distributions, characterized by parameters *a* and *b,* to represent the probability of boarding when purchasing different numbers of tickets, but only the elastic controllability model learns latent elasticity estimates. Specifically, in the ‘controllability model’, three beta distributions represent the expected level of control from purchasing 1, 2, or 3 tickets. Thus, the parameters of these distributions (𝑎_n_, 𝑏_n_, n ∈ {1, 2, 3}) accumulate the number of times each number of tickets led (𝑎_n_) or did not lead (𝑏_n_) to successful boarding. Conversely, in the ‘elastic controllability model’, the beta distributions represent a belief about the maximum achievable level of control (𝑎_control_, 𝑏_control_) coupled with two elasticity estimates that specify the degree to which successful boarding requires purchasing at least one (𝑎_elastic≥1_, 𝑏_elastic≥1_) or specifically two (𝑎_elastic2_, 𝑏_elastic2_) extra tickets. As such, these elasticity estimates quantify how resource investment affects control. The higher they are, the more controllability estimates can be made more precise by knowing how much resources the agent is willing and able to invest (Supplementary Note 1). Figure 4 shows schematically how the two models differ in updating their estimates in response to different types of outcomes. Both models similarly use these estimates to compute an expected value for purchasing 0, 1, 2, or 3 tickets, and accordingly choose between these options (see Methods for model equations and Supplementary Table 3 for all parameter fits, priors, and their interpretation).

**Figure 4.**
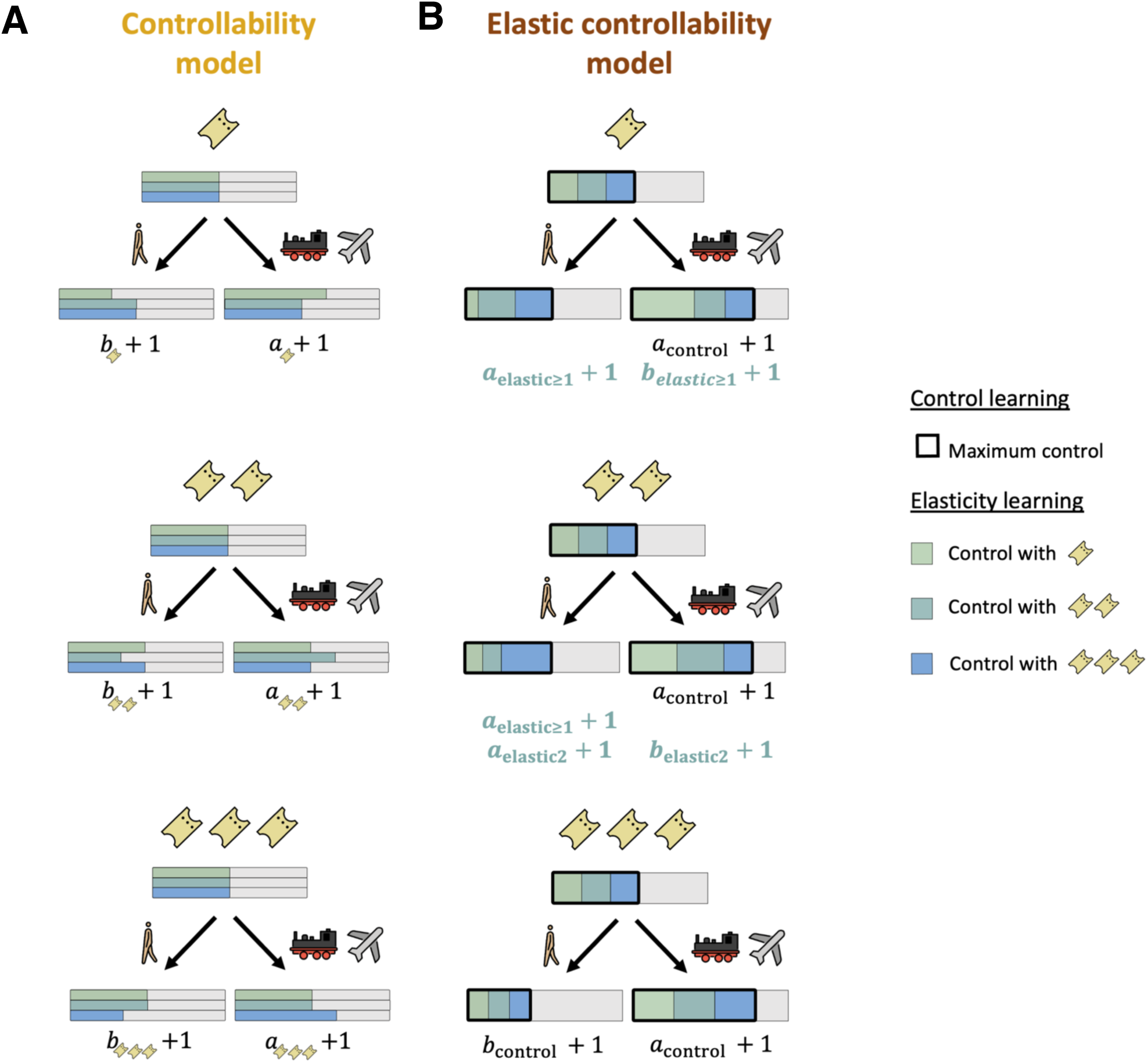
Computational models: Illustration of the learning rules of the *controllability* and *elastic controllability* models. The width of the colored region represents the estimated control, shown as a percentage of absolute control (gray area) along with the update rules based on each outcome. (A) **Controllability model:** This model treats the purchase of 1, 2, and 3 tickets as distinct actions. It accumulates evidence for the effectiveness (𝑎) or lack of effectiveness (𝑏) of each action based solely on whether or not the action led to successful boarding, illustrated by changes in the width of the shaded region corresponding to each ticket amount. (B) **Elastic Controllability model:** Represents beliefs about maximum controllability (black outline) and the degree to which at least one or two extra tickets are necessary to obtain it. These beliefs are used to calculate the expected control when purchasing 1 ticket (inelastic controllability) and the additional control afforded by 2 and 3 tickets (elastic controllability), demonstrated by the bars. The overlapping arrangement of the bars illustrates that, only in the elastic controllability model, maximum controllability is the sum of inelastic and elastic control. The model updates its beliefs as follows: **Top –** Successfully boarding with one ticket provides evidence of high maximum controllability (𝑎_control_ + 1; expanded shaded region) and reduces the perceived need to purchase extra tickets (𝑏_elastic≥1_ + 1; light green expanded). A failure to board does not change estimated maximum controllability, but rather suggests that 1 ticket might not suffice to obtain control (𝑎_elastic≥1_ + 1; light green diminished). As a result, the model’s estimate of control afforded by 1 ticket, as well as on average across ticket options, is reduced. **Middle –** Successfully boarding with 2 tickets provides evidence of high maximum controllability (𝑎_control_ + 1; expanded shaded region), and reduces the perceived need to purchase two extra tickets (𝑏_elastic2_ + 1; light & dark green expanded). Here too, a failure to board does not change estimated maximum controllability, but rather suggests that 2 tickets might not be sufficient, thus reducing the estimated control for 2 tickets and on average (𝑎_elastic2_ + 1; light & dark green diminished). **Bottom** – Successfully boarding with 3 tickets provides evidence of high maximum controllability (𝑎_control_ + 1; expanded shaded region), but offers no insight about whether fewer tickets would suffice to obtain it. Failure to board with 3 tickets provides evidence of low controllability (𝑏_control_ + 1; diminished shaded region).

Examining the predictions of both models, each simulated using the parameter setting that best fit participants’ choices, showed that the elastic-controllability model was better than the controllability model at opting-out in low controllability planets (elastic-controllability model % opt in = 49% ± .01, controllability model % opt in = 63% ± .01; Figure 5C, top panel). Most importantly, this higher level of performance closely matched participants’ behavior (% opt in = 47%, ±.02). Furthermore, the elastic controllability model was also better at not purchasing extra tickets in low-controllability (elastic controllability extra tickets = .49±.01 controllability extra tickets = .65±.01) and inelastic-controllability (elastic controllability extra tickets = .58±.01 controllability extra tickets = .66±.01) planets, and this too was more closely aligned with participants’ behavior (low-controllability extra tickets = 51±.02, inelastic-controllability extra tickets = 53±.02; Figure 5C, bottom panel).

**Figure 5.**
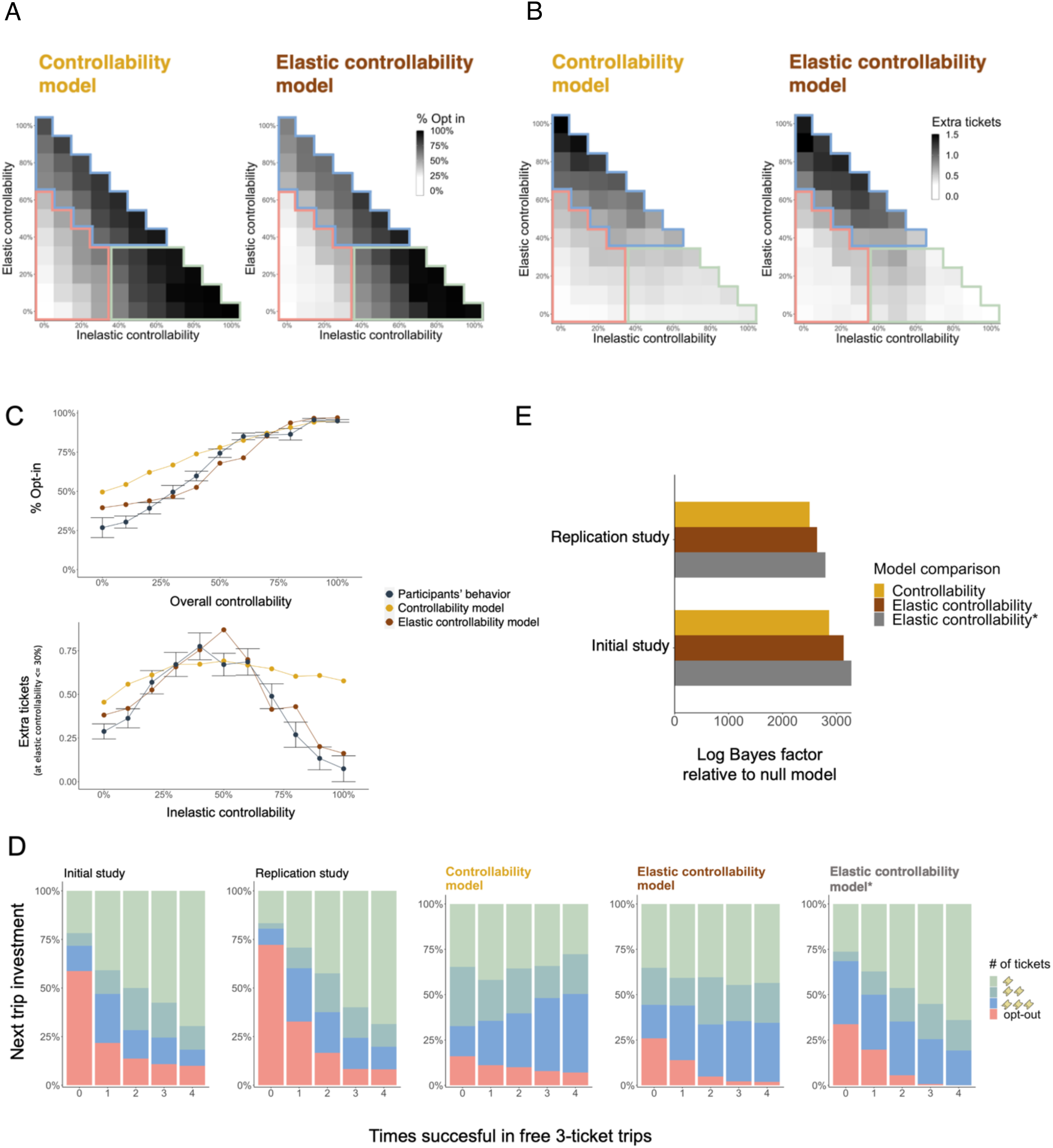
Model simulations: (A) **Opt-in.** Proportion of opt-in by the controllability and elastic controllability models across all planets. Each region is outlined by the color corresponding to the optimal ticket strategy (Blue: 3 tickets, Green: 1 ticket, Red: opt-out). (B) **Extra ticket purchases.** Extra tickets purchased by the controllability and elastic controllability models across all planets. Here too, each region is outlined by the color corresponding to the optimal ticket strategy. (C) **Comparison of opt-in rates and extra ticket purchases made by models and participants. Top -** Proportion of opting-in as a function of overall controllability. **Bottom -** The number of extra tickets purchased for each level of inelastic controllability when elasticity did not warrant purchasing extra tickets (i.e. below 30%). (D) **Resource investment following free 3-ticket trips.** The distribution of resource investment choices made immediately following the free 3-ticket trips, as a function of the outcomes of those trips. Results are shown for the two studies (Initial, Replication), the controllability model, and the original and modified variant of the elastic controllability models. (E) **Model comparison.** Consistency of each model with participant behavior is shown as log Bayes factor compared to a null model wherein participants have distinct preferences among different number of tickets to purchase but no latent controllability estimates.

Beyond these general aspects of task behavior, having participants start on every planet with free 3-ticket trips also enabled us to examine their response to succeeding or failing with 3 tickets, prior to any other experiences on the planet (Figure 5D). As noted above, failure with 3 tickets makes the controllability model favor 1 and 2 tickets over 3 tickets, whereas success with 3 tickets makes it favor 3 tickets over 1 and 2 tickets. Critically, neither the participants nor the simulation of the elastic controllability model showed these effects. Upon failing with 3 tickets, both participants and the elastic model primarily favored to opt out. Conversely, if they succeeded with 3 tickets, they did not become less likely to purchase 1 or 2 tickets.

In fact, failure with 3 tickets even made participants favor 3, over 1 and 2, tickets. This favoring of 3 tickets continued until participants accumulated sufficient evidence about their limited control to opt out (Supplementary Figure 2). Presumably, the initial failures with 3 tickets resulted in an increased uncertainty about whether it is at all possible to control one’s destination. Consequently, participants who nevertheless opted in invested maximum resources to resolve this uncertainty before exploring whether control is elastic. While this behavior directly contradicts the controllability model, it also goes beyond the behavior exhibited by the elastic controllability model. To capture this aspect of participants’ behavior, we modified the elastic controllability model (elastic controllability*) such that it would increase its elasticity estimates when controllability is reduced. Model comparison consistently supported this modification (Initial study: log Bayes Factor = 143 Replication study: log Bayes factor = 155).

More importantly, both the original and modified elastic controllability models explained participants’ behavior better than the controllability model (Initial: log Bayes Factor ≥ 270, Replication: log Bayes Factor ≥ 148; Figure 5E) as well as other theoretically motivated alternatives (see Methods). These results establish that people infer the elasticity of control, and thereby adapt their resource allocation to the environment’s controllability and its elasticity.

### Individual elasticity biases explain misallocation of resources

To determine whether the elastic controllability* model could identify individual biases in elasticity inference, we fitted the model to participants’ choices and thus derived each participant’s beliefs about controllability (𝛾_controllability_) and elasticity (𝛾_elasticity_) prior to visiting any planet. 𝛾_controllability_ is used to initialize the model’s belief about the maximum available control (𝑎_control_, 𝑏_control_), with higher values reflecting stronger prior beliefs that planets are controllable. 𝛾_elasticity_ is used to initialize the model’s elasticity estimates (𝑎_elastic_, 𝑏_elastic_), with higher values reflecting stronger prior beliefs that control is elastic (see Methods). The stronger the belief the greater its influence on choices to opt in (𝛾_controllability_) and purchase extra tickets (𝛾_elasticity_) regardless of observed outcomes. In addition to these two model parameters, we examined four additional parameters that directly influence opting-in and the purchase of extra tickets. These include an inverse temperature parameter (β), which specifies the stochasticity with which expected values affect resource investment choices, and baseline-preference parameters for purchasing 1, 2, or 3 tickets (α1, α2, α3). Across both samples, shared variance among parameters never exceeded 14% (Supplementary Figure 1), and was particularly low for the controllability and elasticity priors (Initial study: 5%, Replication study: 3%). Thus, a tendency to assume the environment is controllable did not imply a tendency to assume that control is elastic, establishing inferences concerning maximum controllability and it elasticity as distinct facets of individual differences.

To clarify how fitted model parameters related to observable behavior, we regressed participants’ opt-in rates and extra ticket purchases on the parameters (Figure 6A). We found that a higher controllability bias (higher 𝛾_controllability_) was associated with more opting in (Initial study *p*<.001, Replication study *p*<.02; logistic regression) and more extra tickets purchased (Initial *p*<.001, Replication *p*<.001; probit regression) on low-controllability, as compared to high-controllability, planets. Thus, a prior assumption that control is likely available (operationalized by 𝛾_controllability_) was reflected in a futile investment of resources in uncontrollable environments.

**Figure 6.**
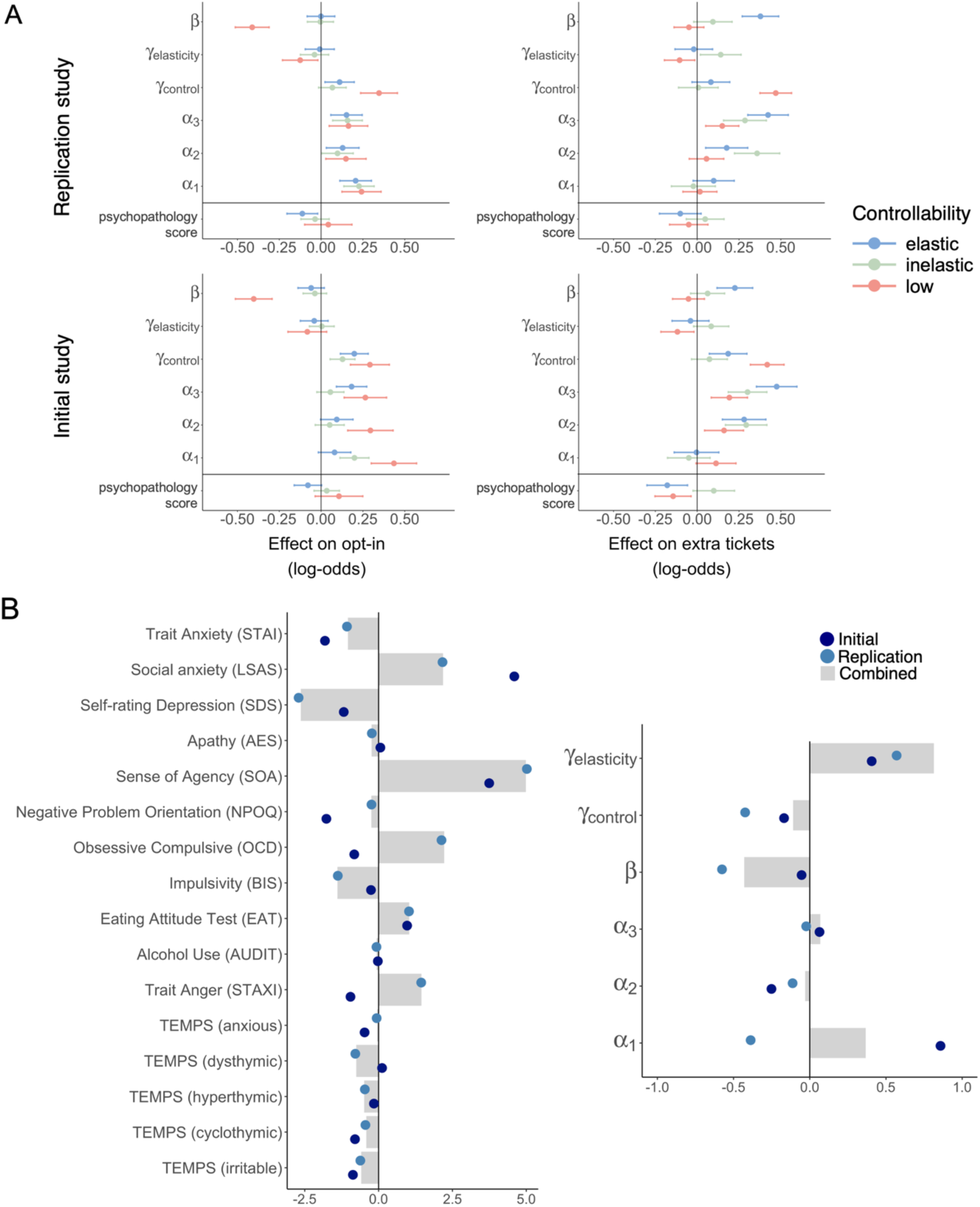
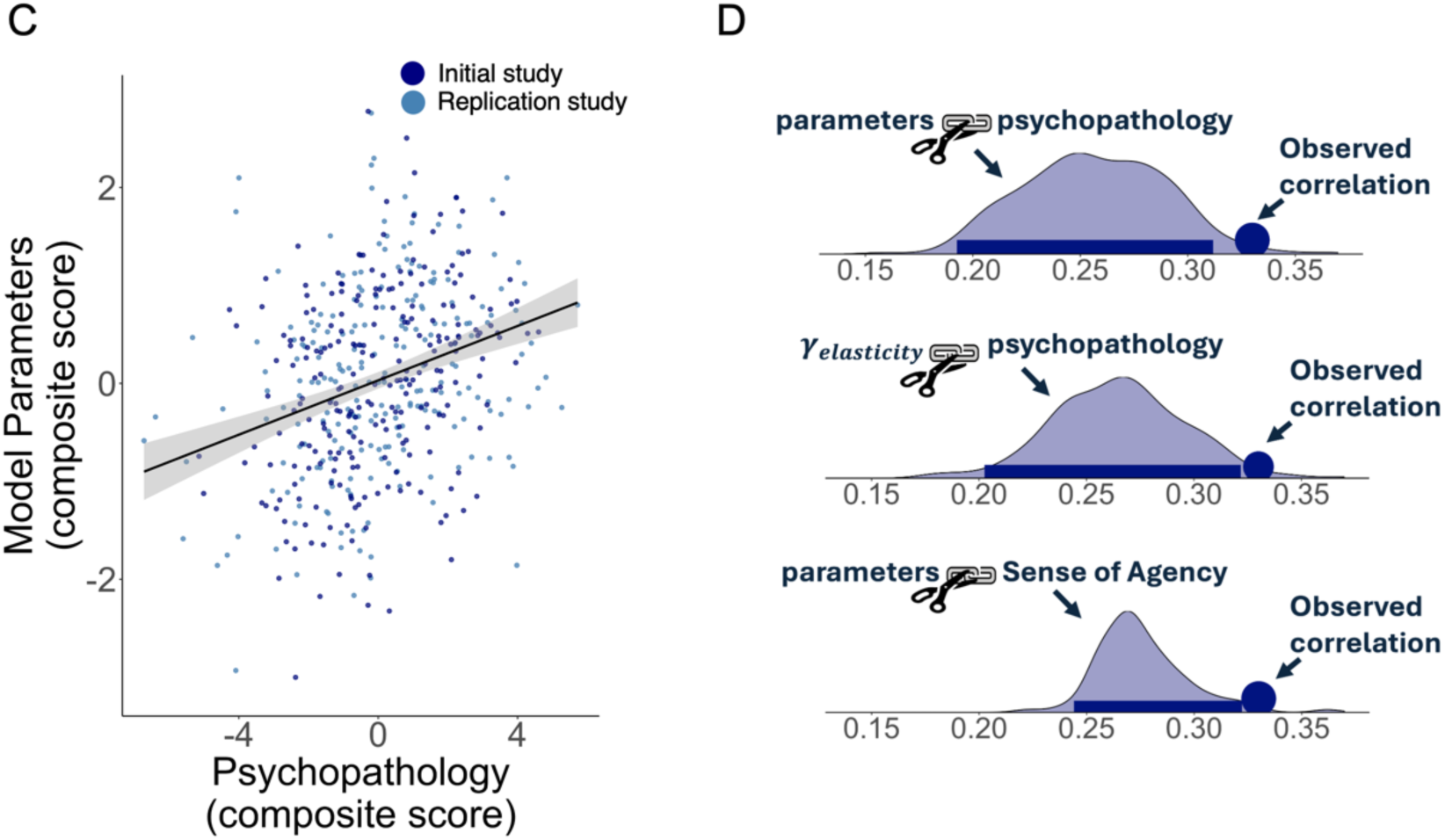
(A) Model parameters and task behavior. Model parameters fitted to each participant’s task behavior were used as predictors of opt-in (left; multiple logistic regressions) and extra ticket purchases (right; multiple probit regressions) separately for low controllability planets where opting out is optimal (red), high elastic controllability planets where purchasing 3 tickets is optimal (blue), and high inelastic controllability planets where purchasing 1 ticket is optimal (green). In a separate analysis, participants’ composite psychopathology scores (see panel C) were similarly used to predict opt-in (left; logistic regression) and extra ticket purchases (right; probit regression). Bars represent regression coefficients and 95% CIs. (B) **Model parameters and psychopathology.** Loadings from a Canonical Correlation Analysis (CCA) between model parameters and self-report psychopathology measures. Self-report measures were scored such that higher values reflect higher levels of psychopathology. Loadings are shown for the Initial (dark blue), Replication (light blue), and combined (gray bars) datasets. The magnitude of the loading reflects the degree to which each measure contributes to the correlation between parameters and self-reports. (C) **Model parameters and psychopathology composite scores.** For each participant, a composite psychopathology score was calculated by multiplying their self-report scores with their CCA-derived loadings, and a composite model parameter score was calculated by multiplying their parameter fits and corresponding loadings. Each dot represents an individual participant. Black line: group-level posterior slope. Gray shading: standard error. (D) **Significance testing.** p-values for the observed CCA correlation (circle) were computed against empirical null distributions (shaded area and bar showing 95% high-density interval). The top null distribution was obtained by shuffling the model parameter fits (top) relative to the psychopathology scores (top), the middle null distribution was obtained by shuffling only the elasticity prior parameter (𝛾_elasticity_) while keeping all other parameters and self-report scores matched (middle), and the bottom null distribution was obtained by shuffling only the Sense of Agency score (SOA) while keeping all other parameters and self-report scores matched (bottom).

In contrast, a higher elasticity bias was associated with the purchasing of more extra tickets on planets with inelastic, as compared to elastic, controllability (Initial study: 𝛽 = .18, 95% CI = [0.03, 0.34], p = .02; Replication study: 𝛽 = .15, 95%CI = [0.02 0.28], p = .02; linear regression). Thus, a prior assumption that control is likely elastic available (operationalized by 𝛾_elasticity_) was primarily reflected in a needless investment of extra resources in environments where control could be obtained more cheaply.

Other parameters showed associations consistent with their roles in the model. Specifically, the inverse temperature parameter (β) was associated with a more optimal allocation of resources, all three baseline preference parameters (α1, α2, α3) were associated with opting in, and α2 and α3, in particular, were also associated with the purchasing of more extra tickets (Figure 6A).

In sum, the model parameters captured meaningful individual differences in how participants allocated their resources across environments, with the controllability parameter primarily explaining variance in resource allocation in uncontrollable environments, and the elasticity parameter primarily explaining variance in resource allocation in environments where control was inelastic.

### Individual elasticity biases are associated with psychopathology

To examine whether the individual biases in controllability and elasticity inference have psychopathological ramifications, we assayed participants on a range of self-report measures of psychopathologies previously linked to a distorted sense of control (see Methods). Examining the direct correlations between model parameters and psychopathology measures (reported in Supplementary Figure 3) does not account for the substantial variance that is typically shared among different forms of psychopathology. For this reason, we instead used a canonical correlation analysis (CCA) to identify particular dimensions within the parameter and psychopathology spaces that most strongly correlate with one another. The CCA loadings indicate the contribution of each parameter of psychopathology score to the overall canonical correlation. We assessed statistical significance of the canonical correlation against a null distribution obtained by permuting the model parameter relative to the psychopathology scores. This showed that the correlation was significant in both the Initial (p=.01) and the Replication (p=.02) studies.

To obtain more robust estimates of the CCA loadings, we next ran a canonical correlation analysis on both datasets combined (Figure 6C, ρ=.33, p=.004). On the side of model parameters, the elasticity prior parameter (𝛾_elasticity_) received the highest loading. By comparison, the controllability prior parameter received a lower loading with the opposite sign. A further permutation test showed that the contribution of the elasticity prior parameter to the parameter-psychopathology correlation was itself significant (*p*=.04; Figure 6D).

Loadings on the side of psychopathology were dominated by an impaired sense of agency (SOA; contribution to canonical correlation: p=.03, Figure 6D, bottom plot), along with obsessive compulsive symptoms (OCD), and social anxiety (LSAS) – all symptoms that have been linked to an impaired sense of control^25–27^. Conversely, depression scores (SDS) received a negative loading, consistent with evidence of a lower willingness to expend one’s resources in depression^23,28–29^. This psychopathological profile also manifested in raw task measures, since, like the elasticity prior parameter, it too was associated with the purchasing of more tickets on planets where controllability was less elastic (Initial: 𝛽 = .19, 95% CI = [. 06, .32], p=.004; Replication: 𝛽 = .13, 95% CI = [. 01, .25], p = .03; linear regression; Figure 6A). Thus, a distinct psychopathological profile involving a distorted sense of agency was associated with a bias in elasticity inference.

## Discussion

Across two pre-registered studies, we demonstrate that humans infer the elasticity of control over their present environment, and this informs their decisions of whether and how to act. Accordingly, we found that a bias to overestimate the elasticity of control leads to a mismanagement of resources, and is associated with elevated psychopathology involving an impaired sense of agency.

Unlike prior work that treated controllability and its elasticity as coupled, our studies dissociated elastic and inelastic controllability, revealing that individuals who typically assume that taking action is worthwhile (overall controllability) do not necessarily assume that investing more resources in taking action is worthwhile (the elasticity of control). In fact, biases regarding controllability and its elasticity were oppositely related to psychopathology, revealing a psychopathological profile that combines underestimation of control with overestimation of the degree to which investing more resources is necessary to obtain control.

The observed elasticity biases across individuals are particularly striking given the simplicity of the experimental task in comparison to real-life scenarios. Unlike the task, where successful boarding directly determined a trip’s outcome, in many real situations many actions may contribute to a single outcome. In this regard, the present work introduces a new kind of credit-assignment problem^30^, which involves identifying not only which actions influenced the outcome but also which of these actions was made more effective by investing more resources in its execution. Moreover, whereas in the task elasticity increased linearly with the number of tickets, resource investment in real life often exhibits non-linear effects. Specifically, further increases in resource investment often result in diminishing returns, and beyond a certain point, even worse outcomes (e.g. hiring too many employees, making the work less efficient). Additionally, real life typically doesn’t offer the streamlined recurrence of homogenized experiences that makes learning easier in experimental tasks, nor are people systematically instructed and trained about elastic and inelastic control in each environment. These complexities introduce substantial additional uncertainty into inferences of elasticity in naturalistic settings, thus allowing more room for prior biases to exert their influences. The elasticity biases observed in the present studies are therefore likely to be amplified in real-life behavior. Future research should examine how these complexities affect judgments about the elasticity of control to better understand how people allocate resources in real-life.

Our finding that elasticity biases are associated with agency-related psychopathologies raises interesting questions for future research on the causal nature of this association. One possibility is that biased elasticity inferences distort one’s sense of agency by leading to maladaptive resource allocation that results in consistently suboptimal outcomes. Another possibility is that people who already have a reduced sense of agency, attempt to compensate for this feeling by investing more resources despite minimal benefit, thus demonstrating an elasticity bias. Importantly, these possibilities are not mutually exclusive. That is, elasticity biases and a distorted sense of agency could reinforce each other in a cyclical manner. Longitudinal studies tracking or manipulating elasticity beliefs while monitoring changes in real-life resource investment and subjective sense of agency, could resolve how elasticity biases and psychopathology influence each other.

Whereas our pre-registered CCA effectively identified associations between task parameters and a psychopathological profile, this analysis method does not directly reveal relationships between individual variables. Auxiliary analyses confirmed significant contributions of both elasticity bias and sense of agency to the observed canonical correlation, but the contribution of other measures remains to be determined by future work. Such work could employ other established measures of agency, including both behavioral indices and subjective self-reports, to better understand how these constructs relate across different contexts and populations. Our operationalization of controllability aligns with previous work that defined controllability as the degree to which actions influence the probability of obtaining a reward^5–6,6^. Other notable works defined controllability as the degree to which actions predict subsequent states, independent of reward^1–3^ Here we adhered to the former definition due to our present focus on resource elasticity, since individuals are unlikely to invest resources to gain control if control does not yield more reward. That said, we followed the latter work in ensuring that controllability is not confounded with predictability^2^, in the sense that in planets with high controllability, outcomes were not necessarily more predictable than in planets with low controllability (See Methods). This enabled us to specifically assess individual inferences concerning controllability and its resource elasticity.

Future research could explore alternative models for implementing elasticity inference that extend beyond our current paradigm. First, further investigation is warranted concerning how uncertainty about controllability and its elasticity interact. In the present study, we minimized individual differences in the estimation of maximum available control by providing participants with three free tickets at their initial visits to each planet. We made this design choice to isolate differences in the estimation of elasticity, as opposed to maximum controllability. To study how these two types of estimations interact, future work could benefit from modifying this aspect of our experimental design.

Second, future models could enable generalization to levels of resource investment not previously experienced. For example, controllability and its elasticity could be jointly estimated via function approximation that considers control as a function of invested resources. Although our implementation of this model did not fit participants’ choices well (see Methods), other modeling assumptions drawn from human functional learning^31^ or experimental designs with continuous action spaces may offer a better test of this idea.

Another interesting possibility is that individual elasticity biases vary across different resource types (e.g., money, time, effort). For instance, a given individual may assume that controllability tends to be highly elastic to money but inelastic to effort. Although the task incorporated multiple resource types (money, time, and attentional effort), the results may differ depending on the type of resources on which the participant focuses. Future studies could explore this possibility by developing tasks that separately manipulate elasticity with respect to different resource types. This would clarify whether elasticity biases are domain-specific or domain-general, and thus elucidate their impact on everyday decision-making.

## Methods

### Pre-registration

Both the Initial and Replication studies were pre-registered prior to data collection. The pre-registrations included hypotheses, sample sizes, exclusion criteria, task design, analysis plans (regressions, CCA), and computational models. Detailed pre-registration documents are available at https://aspredicted.org/Q7M_BHR and https://aspredicted.org/CHW_12H.

### Participants

We used the Prolific online platform (prolific.com) to recruit participants (Initial study N = 264, Replication study N = 250). Participants were selected based on their demonstrated high engagement levels in prior studies, fluency in English, and absence of psychiatric diagnoses. Due to cultural differences in the perception of controllability^32^ participants were recruited from English speaking western countries. The average age of participants across both datasets was 33 years old (SE =.6), and 38% were female.

The sample size was determined using a power analysis as described below. Exclusion criteria were applied to ensure data quality and participant engagement. Specifically, participants were excluded if they demonstrated inattentiveness (defined as more than 8 mistakes on comprehension quizzes during the instruction phase), displayed less than 90% accuracy in selecting the correct vehicle corresponding to its pre-learned destination (mean error rate = 3%, range [0-10%]; indicating a lack of task comprehension), or chose not to purchase any tickets in 90% or more of trials across all experimental conditions (suggesting a lack of engagement with the task). These criteria led to the exclusion of 22 participants (8.3% exclusion rate) from the initial study and 23 participants (8.9% exclusion rate) from the replication study. Participants gave written informed consent before taking part of the study, which was approved by the university’s ethics review board (approval number: 2022-06213). Participants were paid at a rate of 9 euros per hour.

### Experimental Task

Participants were informed about the location of a treasure (150 coins, worth 15¢ of bonus monetary compensation) and could exercise control by boarding the train to reach the house, or the plane to reach the mountain. (see main text and Figure 2). Forgoing or failing to board resulted in walking to the nearest location which happened to be where the treasure was located 20% of the time.

The first ticket (cost = 40 coins) allowed participants to select their desired vehicle, but it only stopped for them at the platform some percentage of the time, corresponding to the inelastic controllability of the planet. Participants could purchase up to 2 additional tickets (cost = 20 coins each) to attempt boarding the moving vehicle after it had left the platform. This improved the probability of boarding in correspondence with the elastic controllability of the planet. When participants purchased a second or third ticket, the chosen vehicle appeared moving from left to right across the screen, and participants attempted to board it by pressing the spacebar when it reached the center of the screen. The precision of each boarding attempt was quantified as the absolute distance of the vehicle from the screen’s center at the time of the press. Unless elastic control was below 15%, boarding attempts with precision in the top 15 percentile relative to the participant’s prior performance automatically succeeded, and unless elastic control was above 85%, those in the bottom 15 percentile automatically failed. For the remaining attempts, the outcome was probabilistically determined such that the overall probability of successfully boarding matched the elasticity of control on that planet. To verify that participants’ pressing ability did not bias their preference for purchasing extra tickets, we ran a mixed probit regression of the number of extra tickets purchased on participants’ average press precision. We found no significant effect (p = .45). When participants purchased multiple tickets, they made all boarding attempts in sequence without intermediate feedback, only learning whether they successfully boarded upon reaching their final destination. This served two purposes. First, to ensure that participants would consistently need to invest not only more money but also more effort and time in planets with high elastic controllability. Second, to ensure that results could potentially generalize to the many real-world situations where the amount of invested effort has to be determined prior to seeing any outcome (e.g., preparing for an exam or a job interview).

At the end of each trip, participants were shown where they arrived, allowing them to infer whether they successfully boarded their vehicle. Since this feedback is only interpretable if participants knew each vehicle’s destination, we only included participants in subsequent analyses if they chose the vehicle corresponding to the destination of the treasure with at least 90% accuracy (Mean 97% ±2%, see first exclusion criteria). The average trial duration was approximately 11 seconds (SE = 0.05), with each additional ticket extending this duration by approximately 0.5 seconds (SE = 0.04), increasing the time investment required from participants.

To allow sufficient learning of each planet’s elasticity and controllability, participants took 30 consecutive trips on each planet. This trip count was determined based on pilot data (N=19) by analyzing when participants’ strategies stabilized. Specifically, we examined the standard deviation in ticket choices over the final 5 trips and found stabilization after around 15 trips (STD = 0.33 compared to .63 in the first 15 trips). Based on this, we provided 15 initial trips per planet for learning, and analyzed ticket choices starting from trip 16 onward. This approach followed our pre-registered plan.

To homogenize learning experience across participants, we gave participants three free tickets for the first five visits to every planet. However, only four of those trips were informative, since on one of every five trips (randomly placed), the treasure was located in the nearby location, making boarding inconsequential for reaching it.

Points accumulated across all planets throughout the session, with participants explicitly motivated to maximize their total points as this directly determined their monetary bonus payment. To ensure that accumulated gains did not lead participants to adopt a simple heuristic strategy of always purchasing 3 tickets, we conducted a mixed probit regression examining whether the number of accumulated coins influenced participants’ decisions to purchase extra tickets. We did not find such an effect (β_coins accumulated_ = .01 p = .87), ruling out the potential strategy shift.

### Planet types

To study elasticity learning across all possible degrees of controllability and elasticity, 66 unique planet conditions were generated by systematically varying inelastic controllability (the probability of the chosen vehicle stopping at the platform if purchasing a single ticket) and elasticity (the added probability of boarding from purchasing 2 extra tickets) from 0 to 100% in 10% increments. The treasure value (150 coins) and ticket costs (40 coins for the first ticket, 20 coins for extra tickets) were chosen so that the optimal strategy, as determined by expected value calculations (*equation 11*), was to purchase 0 tickets on 22 planets (low controllability condition), 1 ticket on a different set of 22 planets (high inelastic controllability condition), and 3 tickets on the remaining 22 planets (high elastic controllability condition). Participants were randomly assigned one planet from each condition, while ensuring they could earn roughly the same total number of coins if they made optimal choices (Initial study= 45±.3, Replication study 46 ± .3). To address potential confounds between controllability and predictability^2^, we designed the experiment such that high-controllability planets (where purchasing 1 or 3 tickets was optimal) were not inherently more predictable than low-controllability planets. We quantified predictability as the entropy of the probability distribution determining whether the participant will or will not arrive at the rewarded location. For 60% of participants, purchasing tickets on high-controllability planets decreased entropy (i.e., increased predictability) compared to low-controllability planets, whereas for the other 40% it increased entropy (i.e., decreased predictability). To increase sensitivity to individual differences, all participants first encountered a planet where purchasing any number of tickets (0, 1, 2, or 3) yielded the same expected value (EV = 30 coins).

### Initial training

Prior to engaging in the task, participants completed 90 practice trials. These practice trials were divided equally across planets with high inelastic controllability, high elastic controllability, and low controllability. To reinforce participants’ understanding of how elasticity and controllability were manifested in each planet, they were informed of the planet type they had visited after every 15 trips. Specifically, for high inelastic controllability planets, the message read: *“Your ride actually would stop for you with 1 ticket! So purchasing extra tickets, since they do cost money, is a WASTE.”* For low controllability planets: *“Doesn’t seem like the vehicle stops for you nor does purchasing extra tickets help.”* Lastly, for high elastic controllability planets: *“Hopefully by now it’s clear that only by purchasing 3 tickets (LOADING AREA) are you consistently successful in catching your ride.”*.

### Power analysis

The sample sizes for both the initial and the replication studies were pre-registered before data collection commenced. For the initial study, we derived the sample size of 264 participants using a power analysis aiming to estimate with high accuracy (credible interval width < 0.1) the likelihood that participants will opt-in for any combination of elastic and inelastic control (66 total combinations). For this purpose, we used Kruschke’s (2014) procedure for Bayesian Power Analysis^33^. Based on pilot data (N=19), we fitted a Bayesian Beta Binomial model to the proportion of opt-ins in the planet with no optimal strategy (and therefore the highest variability in behavior would be expected), and found that 12 participants were required for the credible interval of the binomial proportion parameter to be lower than 0.1. Since each participant completes 3 combinations of elastic and inelastic control, we multiplied the required sample size by the total number of planets in each condition (22) which results in an estimate of n=264.

To examine whether this sample size is sufficient to detect significant effects of elastic and inelastic controllability on opting in (Figure 3A,C), and the purchasing of extra tickets (Figure 3B,C), we used a bootstrap procedure whereby we sampled 264 participants from the pilot data and ran the regression analyses on them. We repeated this procedure for 50 iterations, each time recording 𝛽_elastic_, 𝛽_inelastic_, and their p-values. For analysis 1, we found that both 𝛽_elastic_ 𝑎𝑛𝑑 _inelastic_ were significantly above 0 in all iterations, and for analysis 2, 𝛽_elastic_ was significantly above 0 in all iterations, thereby giving us >98% power to detect all of our effects of interest. Confidence intervals were derived from a binomial test in which a positive significant coefficient counted as a success.

For the replication study, we derived the sample size of 250 participants using a power analysis aiming for 95% power to detect a significant (p <.05) canonical correlation between the model-derived parameters and questionnaire scores (Figure 6B). For this purpose, we sampled with replacement from the initial dataset (n=264) which had showed a significant canonical correlation (p=.01; Figure 6B), and calculated the p-value of a CCA in the subsample. We repeated this procedure for 100 iterations, each time recording the calculated p-value. To ensure our sample size was sufficient to detect significant effects in the regression models predicting opt-in and extra ticket purchases from the composite psychopathology score with over 95% power (Figure 6C) we conducted another bootstrap analysis. We sampled 250 participants with replacement each time calculating the composite score and ran the regression analyses on that score, repeating this procedure 100 times. We found that 𝛽_composite_ was significant (p<.05) in 95% of iterations (95% power).

### Regression analyses

Mixed model regression analyses were performed using the “ordinal” package which fits cumulative mixed models via Laplace approximations^34^. The model predicts the likelihood of purchasing tickets (logistic model) and how many extra tickets are purchased (probit model) on the last 15 trips on each planet. Predictors included fixed and random effects for elastic and inelastic controllability and fixed and random intercepts per participant and planet number, with the latter included to account for potential learning or fatigue effects. Statistical significance was assessed using the Wald Z Test^35^ with normality assessed using the Shapiro-Wilk Test^36^.

### Computational Modeling

To assess whether participants inferred elasticity or merely learned the control afforded by each course of action separately, we fit two competing reinforcement-learning models (‘*controllability*’ and ‘*elastic controllability’*). Both models use beta distributions defined by parameters (*a, b*), to form controllability beliefs for each ticket quantity n (n ∈ {1, 2, 3}), but only the elastic controllability model leverages the dependencies that exist between different levels of resource investment. Below, we expand on the computations underlying both models.

### Controllability model

The controllability model accumulates the number of times that each ticket quantity *n* led to successful or unsuccessful boarding:

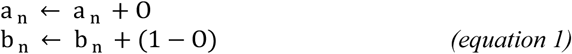

Where O=1 if boarding was successful and O=0 if not. The probability of boarding with 1,2 or 3 tickets on planet s is thus simply the expected value of that particular beta distribution:

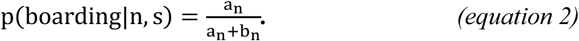

Here and in all subsequent models, learning only takes place when the treasure is not located on the nearby planet, which makes it possible to identify boarding success.

### Elastic controllability model

The *elastic controllability model* explains ticket choices based on latent beliefs about the controllability and elasticity of each planet (see Figure 4). These beliefs are represented by three beta distributions, each defined by two parameters (𝑎, 𝑏). The expected value of one beta distribution (defined by 𝑎_control_, 𝑏_control_) represents the belief that boarding is possible (controllability) with any number of tickets whereas the expected value of the other two beta distributions represent the belief that successful boarding is added by purchasing at least one (𝑎_elastic≥1_, 𝑏_elastic2_) or specifically two (𝑎_elastic2_, 𝑏_elastic2_), extra tickets (elasticity).

Thus, the parameter 𝑎_control_ accumulates the number of successful boarding attempts with any number of tickets, whereas 𝑏_control_ accumulates the number of failed boarding attempts despite purchasing the maximum number of tickets (3 tickets):

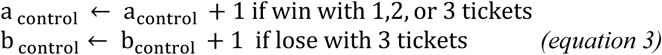

A failure to board with fewer tickets doesn’t reduce the maximum controllability estimate but instead suggests that more tickets or the maximum number of tickets may be needed to obtain control:

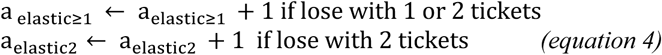

Conversely, successfully boarding with fewer tickets is evidence that obtaining controllability does not require maximal or any extra tickets:

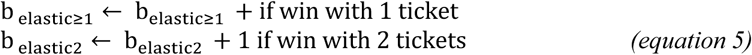

To account for our finding that failure with 3 tickets made participants favor 3, over 1 and 2, tickets, we introduced a modified elastic controllability* model, wherein purchasing extra tickets is also favored upon receiving evidence of low controllability (loss with 3 tickets). This effect was modulated by a free parameter 𝜅 which reflects a tendency to prioritize resolving uncertainty about whether control is at all possible by investing maximum resources:

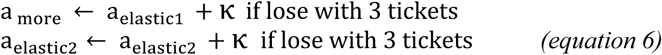

The estimated controllability and elasticity, as characterized by the expected value of the three beta distributions (𝐶_total control_, 𝐶_more_, 𝐶_full_), is used to derive the probability of boarding the selected vehicle with 1, 2, or 3 tickets. Thus, the estimated probability of boarding with 1 ticket is an estimate of inelastic control, that is, the percentage of the total control 𝐶_total control_ that does not require investing more resources:

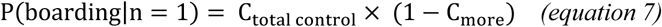

For 2 tickets, the added probability of boarding is an estimate of what percentage of total control is elastic (C_more_) but does not require investing maximal resources (1 − C_full_):

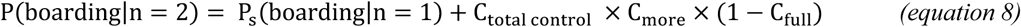

Finally, since 3 tickets is the maximum resource investment, the estimated probability of boarding with 3 tickets is equivalent to an estimate of total attainable control.

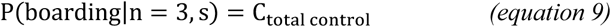

### Models’ resource investment

Because participants win 20% of the time even when they fail to board their selected ride, we calculate the probability of arriving at the treasure with *n* tickets as:

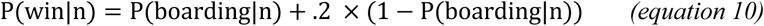

Then, taking the possible reward (R = 150) and ticket costs (C = {40,20,20}) into account, the expected value for purchasing n tickets (n = {1,2,3}) tickets is:

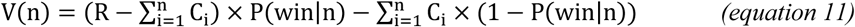

One exception to this rule is the expected value of opting out, V(0), which is similar across all planets, and equals the potential reward (150) times the probability that walking will lead to the reward (0.2).

The probability that a participant will purchase 0, 1, 2, or 3 tickets is calculated using the SoftMax function such that:

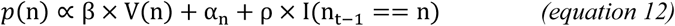

where β is an inverse temperature parameter, α_1–3_ are parameters that represent participants’ fixed tendencies to purchase 1, 2, or 3 tickets respectively, and ρ is a perseveration parameter that represents participant tendency to purchase the same number of tickets as in the last trial (n_t–1_). Α_O_ is set to 0 to maintain parameter identifiability.

We compared the aforementioned models to a null model wherein participants have distinct preferences among different number of tickets (α_1–3_) and perseveration (ρ) but no latent controllability estimates (𝛽 = 0 in equation 12). We also evaluated variants of both models that replaced the SoftMax-based exploration (governed by inverse temperature β) with a decaying epsilon-greedy approach. This epsilon-greedy variant, which begins with a fixed exploration rate that decreases with the number of trials^59^, performed significantly worse at explaining participants’ choices compared to the SoftMax-based models.

### Prior controllability and elasticity beliefs

To capture prior biases that planets are controllable and elastic, we introduced parameters 𝛾_controllability_ and 𝛾_elasticity_, each computed by multiplying the direction (λ – 0.5) and strength (ɛ) of individuals’ prior belief. λ_controllability_ and λ_elasticity_ range between 0 and 1, with values above 0.5 indicating a bias towards high controllability or elasticity, and values below 0.5 indicating a bias towards low controllability or elasticity. ɛ_controllability_ and ɛ_elasticity_ are positively valued parameters capturing confidence in the bias. Parameter recovery analyses confirmed both good recoverability (see S2 Table) and low confusion between bias direction and strength ^(ɛ^controllability^→λ^controllability ^= −.07^, λcontrollability^→ɛ^controllability ^=^ −.16, ɛ_elasticity_→λ_elasticity_ = .15, λ_elasticity_→ɛ_elasticity_ = .04), ensuring that observed biases and their relation to psychopathology do not merely reflect slower learning (Supplementary Figure 4), which can result from changes in bias strength but not direction.

For each new planet participants encountered, these parameters were used to initialize the beta distributions representing participants’ beliefs:

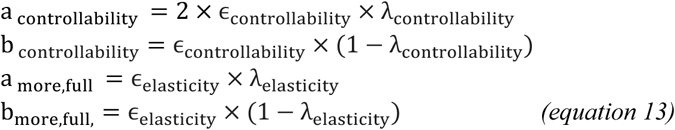

𝑎 _controllability_ is multiplied by 2 so that in the absence of a bias, it would have equal preference for purchasing 0 to 3 tickets (this requires that C _controllability_ = 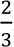, C _more,full_ = 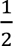).

For consistency, in the ‘controllability model’, the controllability estimates for each ticket quantity are initialized for each ticket quantity n, where n={1,2,3}, as:

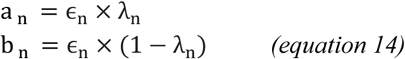

Here, to ensure no preference between different ticket quantities in the absence of a bias, 𝑎_1_ is multiplied by 2 and 𝑎_2_ is multiplied by 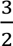.

### Testing alternative models

To further validate our elastic controllability model, we tested it against several theoretically motivated alternatives that could potentially explain participants’ behavior.

To ascertain that participants were truly learning latent estimates of controllability rather than simpler associations, we conducted two complementary analyses.

First, we implemented a simple Q-learning model that directly maps ticket quantities to expected values based on reward prediction errors, without representing latent controllability. This associative model performed substantially worse than even our simple controllability model (log Bayes Factor ≥ 1854 on the combined datasets). Second, we fitted a variant of the elastic controllability model that compared learning from control-related versus chance outcomes via separate parameters (instead of assuming no learning from chance outcomes). Chance outcomes were observed by participants in the 20% of trials where reward and control were decoupled, in the sense that participants reached the treasure regardless of whether they boarded their vehicle of choice. Results showed that participants learned considerably more from control-related, as compared to chance, outcomes (mean learning ratio=1.90, CI= [1.83, 1.97]). Together, these analyses show that participants were forming latent controllability estimates rather than direct action-outcome associations.

Having established that participants indeed learn latent controllability estimates, we next examined our model’s interpretation of the κ parameter. Our model accounts for the observation that failures with 3 tickets made participants more likely to purchase 3 tickets when they opted in. To test whether this behavior resulted from participants not accepting that the planet is uncontrollable, we implemented a variant of the model where κ not only increases the elasticity estimate but also reduces the controllability update (using 𝛽_control_+(1-𝜅) instead of 𝛽_control_+1) after failures with 3 tickets. However, implementing this coupling diminished the model’s fit to the data (log Bayes Factor ≥ 48), as compared to allowing both effects to occur independently, indicating that the increase in 3 ticket purchases upon failing with 3 tickets did not result from participants not accepting that controllability is in fact low. Thus, we maintained our original interpretation that failure with 3 tickets increases uncertainty about whether control is possible at all, leading participants who continue to opt in to invest maximum resources to resolve this uncertainty.

Having validated our interpretation of κ, we next examined whether participant behavior would be better characterized as a continuous function approximation rather than the discrete inferences in our model. To test this, we implemented a Bayesian model where participants continuously estimate both controllability and its elasticity as a mixture of three archetypal functions mapping ticket quantities to success probabilities. The flat function provides no control regardless of how many tickets are purchased (corresponding to low controllability). The step function provides full control as long as at least one ticket is purchased (inelastic controllability). The linear function linearly increases control with the number of extra tickets (i.e., 0%, 50%, and 100% control for 1, 2, and 3 tickets, respectively; elastic controllability). The model computes the likelihood that each of the functions produced each new observation, and accordingly updates its beliefs. Using these beliefs, the model estimates the probability of success for purchasing each number of tickets, allowing participants to weigh expected control against increasing ticket costs. Despite its theoretical advantages for generalization to different ticket quantities, this continuous function approximation model performed significantly worse than the elastic controllability model (log Bayes Factor > 4100 on combined datasets), suggesting that participants did not assume that control increases linearly with resource investment.

### Model fitting

Reinforcement-learning models were fitted to all choices made by participants via an expectation maximization approach used in previous work^37–39^. We generated random parameter settings drawn from predefined group-level distributions and calculated the likelihood of observing the participants’ choices given each parameter setting. We then approximated the posterior estimates of the group-level prior distributions for each parameter by resampling the parameter values, weighted by their respective likelihoods, and refitted the data based on the updated priors. This iterative process of resampling and refitting continued until no further improvement to model fit. To determine the best-fitting parameters for each individual participant, we computed a weighted average of the final batch of parameter settings, where each setting was weighted by the likelihood it assigned to the individual participant’s choices.

Model parameter 𝛽 was initialized by sampling from a Gamma distribution (*k* = 1, *θ* = 1), 𝜆 was initialized by sampling from a Beta distribution (*α* = 1 *and β* = 1), 𝜖 was initialized by sampling from a Lognormal distribution (*μ* = 0, *σ* = 1)., and all other parameters (𝛼_1–3_, 𝜌, 𝜅) were initialized by sampling from a Normal distribution (*μ* = 0, *σ* = 1).

### Validation of model fitting procedure

To validate the model fitting procedure, we simulated data using each model and using the model comparison procedure to recover the correct model (Supplementary Table 1). Furthermore, we validated our parameter fits by simulating data using the best fitting parameters for each parameter and then recovering those parameters. The correlations for the parameters of interest were all equal or greater than *r* = .74 and at least .63 for all other parameters (see Supplementary Table 2).

### Model comparison

To compare between models, in terms of how well each model accounted for participants’ choices, we employed the integrated Bayesian Information Criterion (iBIC)^40–41^. First, we estimated the evidence for each model (L) by calculating the mean likelihood of the model given 200,000 random parameter settings sampled from the fitted group-level prior distributions. Subsequently, we computed the iBIC by penalizing the model evidence to account for model complexity, following this formula: iBIC = 2 ln L + k ln n, where k represents the number of fitted parameters, and n is the number of subject choices used to compute the likelihood. Lower iBIC values indicate a more parsimonious and better-fitting model.

### Model simulations

To examine the predictions of the ‘controllability’ and ‘elastic controllability’ models, we simulated choices using the best-fitting parameter settings obtained from participants’ actual choices. For generality, planets were randomized in the simulation as they were in the actual experiment, such that a simulated participant did not visit precisely the same planets as the actual participant. Furthermore, to avoid discrepancies between models, each planet was repeated the same number of times for both models. 2000 simulations were performed with each model.

### Individual differences

We examined individual biases in elasticity inference using participants’ best-fitting parameters for beliefs about controllability (𝛾_controllability_) and elasticity (𝛾_elasticity_). In addition to these focal parameters, we also examined four other parameters that directly influence opt-in decisions and extra ticket purchases: an inverse temperature/noise parameter (β) and three baseline preference parameters (α1, α2, α3).

We then regressed participants’ actual opt-in (logistic regression) and extra ticket purchases (probit regression) in each of the 3 controllability conditions on participants’ parameters. Regressions were performed in the lme4 package in R using Rstudio^42^. Statistical significance was assessed using the Wald Z Test^35^ with normality assessed using the Shapiro-Wilk Test^36^.

### Self-report measures

To assess trait-level agency beliefs and related psychopathologies, we administered the Sense of Agency Scale (SOA)^43^. Additionally, we measured dysfunctional attitudes toward problem-solving via the Negative Problem Orientation Questionnaire (NPOQ) ^44^, and temperamental differences (five domains: hyperthymic, dysthymic, cyclothymic, irritable, and anxious) via the Temperament Evaluation Questionnaire (TEMPS-A)^45–46^.

Additionally, we incorporated a selection of questionnaires used by Gillan et al. (2016)^47^, which assess three factors—Anxious-depression, Compulsive behavior and intrusive thought, and Social withdrawal—linked to goal-directed control deficits. These include the Apathy Evaluation Scale^48^ (AES), Eating Attitudes Test^49^ (EAT), Alcohol Use Disorders Identification Test^50^ (AUDIT), Barratt Impulsiveness Scale^51^ (BIS), Liebowitz Social Anxiety Scale^52^ (LSAS), Self-Rating Depression Scale^53^ (SDS), Trait Subscale of the State-Trait Anxiety Inventory for Adults^54^ (STAI; Spielberger, 1983), and the revised Obsessive-Compulsive Inventory^55^ (OCI-R). We utilized abbreviated versions of these questionnaires following the methodology of Wise and Dolan (2020)^56^, who refined the questionnaires using a classifier to retain the items most contributory to variance in the three factors without significant information loss.

As preregistered, we aggregated the scores of the selected items from each questionnaire independently for inclusion in our CCA. Questionnaires were scored following the corresponding scoring guide, but in such a way that larger scores always indicate greater psychopathology. Based on best practice recommendations^57–58^, participants were screened for inattentive responding with infrequency items (‘I competed in the 1917 Summer Olympic Games’) which resulted in the removal of 11 participants (Initial: 6, Replication: 5).

Directly using the scores from questionnaires of varying lengths in the CCA can give undue influence to short questionnaires, whose scores are likely more noisy. To mitigate this problem, we adjusted the standard deviation of each questionnaire’s score proportionally to the square root of its number of items before conducting the CCA (in accordance with the formula for standard error). After performing the CCA on these weighted scores and normalized model parameters, the resulting canonical loadings were transformed back to their original standardized scale, ensuring that the reported loadings accurately reflected the original, unweighted scale of the questionnaire scores.

### Canonical Correlation Analysis (CCA)

Canonical correlation analysis was implemented using the robust CCA method based on projection pursuit (’ccAPP’ package;)^59^. We searched for the single canonical dimension that maximizes the Spearman correlation between the model parameters and psychopathology scores. Significance testing was conducted using a two-tailed permutation test with 1,000 iterations. In each iteration, the model parameters were shuffled relative to the psychopathology scores, and the canonical correlation was recalculated. The observed canonical correlation was then compared to this null distribution to obtain a p-value. To evaluate the contribution of the elasticity prior parameter (𝛾_elasticity_) to the parameter-psychopathology correlation, we performed an additional permutation test. In this test, only the elasticity bias estimates (𝛾_elasticity_) were shuffled, while all other model parameters and self-report scores remained constant.

To assess how the identified psychopathological profile manifests in raw task measures, we calculated a composite psychopathology score for each participant. This score was derived by multiplying individual participant scores by CCA-derived loadings, representing their position on the psychopathology spectrum. We then used this composite score as a predictor (𝛽_𝑐omposite_) in a linear regression model to forecast the difference in average opt-in and extra ticket behavior between conditions where controllability was high and inelastic versus high and elastic.

## Supplementary Information

### Supplementary Note 1: Higher elasticity does not imply higher controllability

Here, we show using two established definitions of controllability that whether an environment is more or less elastic does not determine whether it is more or less controllable.

***Definition 1, reward-based controllability***^15^: If control is defined as the fraction of available reward that is controllably achievable, and we assume all participants are in principle willing and able to invest 3 tickets, controllability can be computed in the present task as:

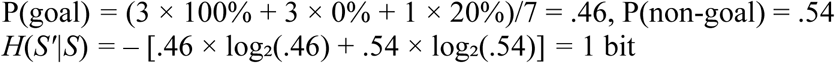

where P(S^′^ = goal ∣ 𝑆, 𝐴, 𝐶) is the probability of reaching the treasure from present state 𝑆 when taking action *A* and investing *C* resources in executing the action. In any of the task environments, the probability of reaching the goal is maximized by purchasing 3 tickets (𝐶 = 3) and choosing the vehicle that leads to the goal (𝐴 = correct vehicle). Conversely, the probability of reaching the goal is minimized by purchasing 3 tickets (𝐶 = 3) and choosing the vehicle that does not lead to the goal (𝐴 = wrong vehicle). This calculation is thus entirely independent of elasticity, since it only considers what would be achieved by maximal resource investment, whereas elasticity consists of the reduction in controllability that would arise if the maximal available 𝐶 is reduced. Consequently, any environment where the maximum available control is higher yet varies less with resource investment would be more controllable and less elastic.

Note that if we also account for ticket costs in calculating reward, this will only reduce the fraction of achievable reward and thus the calculated control in elastic environments.

***Definition 2, information-theoretic controllability***^1–2^: Here controllability is defined as the reduction in outcome entropy due to knowing which action is taken:

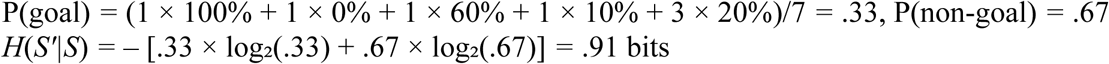

where *H*(*S’|S*) is the conditional entropy of the distribution of outcomes *S’* given the present state 𝑆, and *H*(*S’|S, A, C*) is the conditional entropy of the outcome given the present state, action, and resource investment.

To compare controllability, we consider two environments with the same maximum control:

- Inelastic environment: If the correct vehicle is chosen, there is a 100% chance of reaching the goal state with 1, 2, or 3 tickets. Thus, out of 7 possible action-resource investment combinations, three deterministically lead to the goal state (≥1 tickets and correct vehicle choice), three never lead to it (≥1 tickets and wrong vehicle choice), and one (0 tickets) leads to it 20% of the time (since walking leads to the treasure on 20% of trials).
- Elastic Environment: If the correct vehicle is chosen, the probability of boarding it is 0% with 1 ticket, 50% with 2 tickets, and 100% with 3 tickets. Thus, out of 7 possible action-resource investment combinations, one deterministically leads to the goal state (3 tickets and correct vehicle choice), one never leads to it (3 tickets and wrong vehicle choice), one leads to it 60% of the time (2 tickets and correct vehicle choice: 50% boarding + 50% × 20% when failing to board), one leads to it 10% of time (2 ticket and wrong vehicle choice), and three lead to it 20% of time (0-1 tickets).

Here we assume a uniform prior over actions, which renders the information-theoretic definition of controllability equal to another definition termed ‘instrumental divergence’^60,61^. We note that changing the uniform prior assumption would change the results for the two environments, but that would not change the general conclusion that there can be environments that are more controllable yet less elastic.

#### Step 1: Calculating *H*(*S’|S*)

For the inelastic environment:

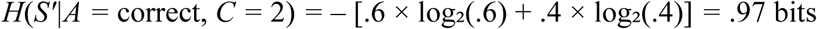

For the elastic environment:

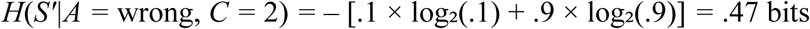

#### Step 2: Calculating *H(S’|S, A, C)*

Inelastic environment: Six action-resource investment combinations have deterministic outcomes entailing zero entropy, whereas investing 0 tickets has a probabilistic outcome (20%). The entropy for 0 tickets is: *H*(*S’|C* = 0) = -[.2 × log₂(.2) + 0.8 × log₂(.8)] = .72 bits. Since this action-resource investment combination is chosen with probability 1/7, the total conditional entropy is approximately .10 bits

Elastic environment: 2 actions have deterministic outcomes (3 tickets with correct/wrong vehicle), whereas the other 5 actions have probabilistic outcomes:

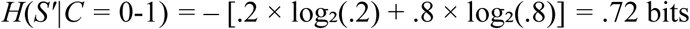

Thus the total conditional entropy of the elastic environment is: *H*(*S’|S, A, C*) *=* (1/7) × .97 + (1/7) × .47 + (3/7) × .72 *=* .52 bits

#### Step 3: Calculating *I(S’ | A, S)*

Inelastic environment: I(*S’*; *A, C | S*) *= H*(*S’|S*) – *H*(*S’|S, A, C*) *=* 1 – 0.1 *=* .9 bits

Elastic environment: I(*S’*; *A, C | S*) *= H*(*S’|S*) – *H*(*S’|S, A, C*) *=* .91 – .52 *=* .39 bits

Thus, the inelastic environment offers higher information-theoretic controllability (.9 bits) compared to the elastic environment (.39 bits).

Of note, even if each combination of cost and success/failure to reach the goal is defined as a distinct outcome, then information-theoretic controllability is higher for the inelastic (2.81 bits) than for the elastic (2.30 bits) environment.

---

**Supplementary Table 1.** Model validation. 10 full experimental datasets were simulated using each model. Rows indicate the model used to simulate data and columns indicate the model recovered from the data using the model comparison procedure.

|  | Controllability | Elastic controllability | Null |
| --- | --- | --- | --- |
| Controllability | 10 | 0 | 0 |
| Elastic controllability | 0 | 10 | 0 |
| Null | 0 | 0 | 10 |

**Supplementary Table 2.** Parameter Recovery. We validated our model fitting procedure by simulating data using the best fitting parameters for each subject and then recovering those parameters. Our correlation between simulated and recovered parameters was at least .74 for all parameters of interest that capture the effects of the experimental conditions, and at least .59 for all other parameters.

| Parameter | Correlation between Simulated and Recovered Parameters |
| --- | --- |
| $\beta$ | .91 |
| $\lambda_{\text{controllability}}$ | .72 |
| $\epsilon_{\text{controllability}}$ | .63 |
| $\gamma_{\text{controllability}}$ | .77 |
| $\lambda_{\text{elasticity}}$ | .81 |
| $\epsilon_{\text{elasticity}}$ | .59 |
| $\gamma_{\text{elasticity}}$ | .74 |
| $\alpha_1$ | .82 |
| $\alpha_2$ | .81 |
| $\alpha_3$ | .81 |
| $\rho$ | .89 |
| $\kappa$ | .63 |

**Supplementary Table 3.** Model parameters. . The table presents means, standard deviations, prior distributions, and interpretations for each parameter of the three computational models. Effective controllability and elasticity biases are computed as 𝛾 = (𝜆 – 0.5)× 𝜖. 𝜅 is only included in the modified elastic controllability* model.

| Elastic controllability model |  |  |  |  |
| --- | --- | --- | --- | --- |
| Parameter | Mean | SD | Prior | Interpretation |
| $\beta$ | 0.07 | 0.02 | $\beta(1,1)$ | SoftMax decision noise |
| $\lambda_{controllability}$ | .69 | .17 | $\beta(1,1)$ | Controllability bias |
| $\epsilon_{controllability}$ | 19.9 | 20 | $\exp(N(0,1))$ | Controllability bias strength |
| $\lambda_{elasticity}$ | .16 | .14 | $\beta(1,1)$ | Elasticity bias |
| $\epsilon_{elasticity}$ | .61 | .20 | $\exp(N(0,1))$ | Elasticity bias strength |
| $\rho$ | 2.2 | 0.66 | $N(0,1)$ | Choice perseveration |
| $\alpha_1$ | 1.3 | 0.28 | $N(0,1)$ | Fixed preference for purchasing 1 ticket |
| $\alpha_2$ | 0.63 | 0.37 | $N(0,1)$ | Fixed preference for purchasing 2 tickets |
| $\alpha_3$ | -0.74 | 0.37 | $N(0,1)$ | Fixed preference for purchasing 3 tickets |
| $\kappa$ | 0.13 | 0.11 | $N(0,1)$ | Prioritize resolving max controllability upon failing with 3 tickets |
| **Controllability model** |  |  |  |  |
| **Parameter** | **Mean** | **SD** | **Prior** | **Interpretation** |
| $\beta$ | 0.06 | 0.01 | $\beta(1,1)$ | SoftMax decision noise |
| $\lambda_1$ ticket | .65 | .20 | $\beta(1,1)$ | 1 ticket bias |
| $\epsilon_1$ ticket | .36 | .42 | $\exp(N(0,1))$ | 1 ticket bias strength concentration |
| $\lambda_2$ tickets | .52 | .12 | $\beta(1,1)$ | 2 ticket bias |
| $\epsilon_2$ tickets | .88 | .52 | $\exp(N(0,1))$ | 2 ticket bias strengthconcentration |
| $\lambda_3$ tickets | .11 | .04 | $\beta(1,1)$ | 3 ticket bias |
| $\epsilon_3$ tickets | 7.97 | .91 | $\exp(N(0,1))$ | 3 ticket bias strengthconcentration |
| $\rho$ | 2.03 | 0.64 | $N(0,1)$ | Choice perseveration |
| $\alpha_1$ | .25 | 0.58 | $N(0,1)$ | Fixed preference for purchasing 1 ticket |
| $\alpha_2$ | 0.47 | 0.57 | $N(0,1)$ | Fixed preference for purchasing 2 tickets |
| $\alpha_3$ | 1.78 | 0.59 | $N(0,1)$ | Fixed preference for purchasing 3 tickets |
| **Null model** |  |  |  |  |
| **Parameter** | **Mean** | **SD** | **Prior** | **Interpretation** |
| $\rho$ | 2.34 | 0.59 | $N(0,1)$ | Choice perseveration |
| $\alpha_1$ | .19 | 0.35 | $N(0,1)$ | Fixed preference for purchasing 1 ticket |
| $\alpha_2$ | .05 | 0.41 | $N(0,1)$ | Fixed preference for purchasing 2 tickets |
| $\alpha_3$ | -.33 | .75 | $N(0,1)$ | Fixed preference for purchasing 3 tickets |

**Supplementary Figure 1.**
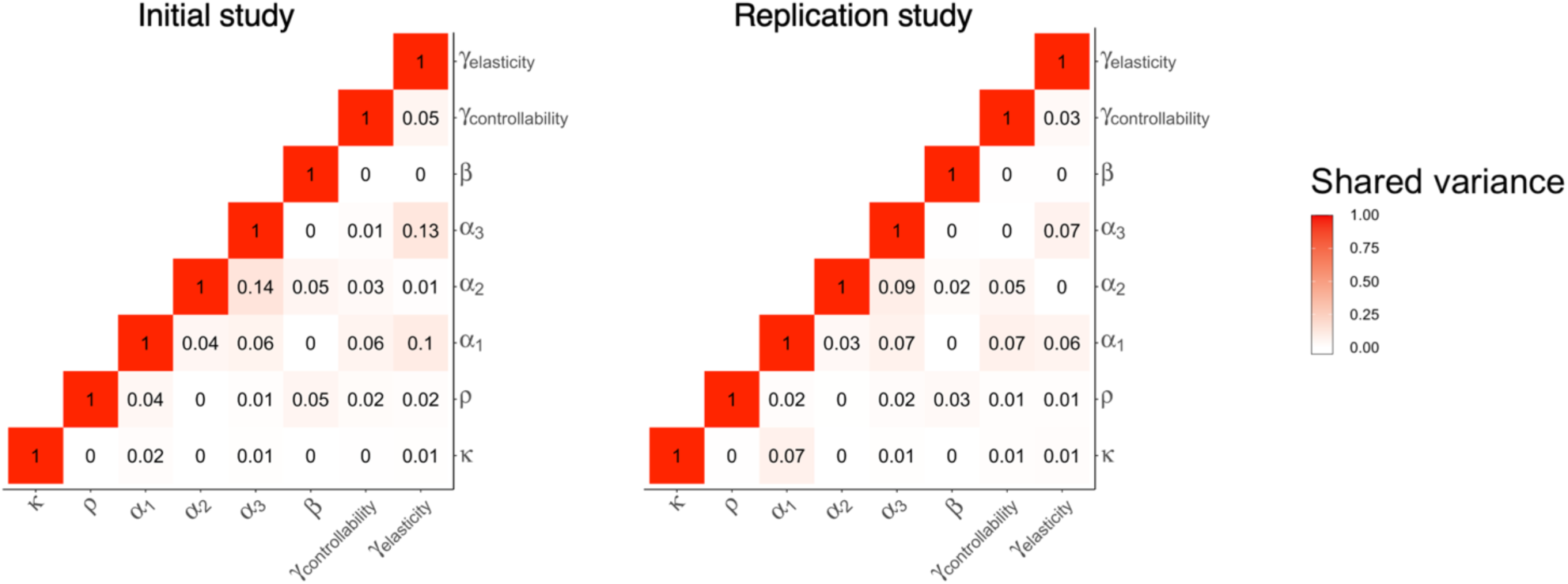
Shared variance between elastic controllability* model parameters. Heatmaps display squared Pearson correlations between parameter pairs for initial (left) and replication (right) studies. Color intensity indicates shared variance magnitude (0-1). Parameters: 𝛾_controllability_, 𝛾_elasticity_, β, 𝑎_1–3_, ρ, κ. Diagonal elements=1; off-diagonal elements reveal parameter interdependencies.

**Supplementary Figure 2.**
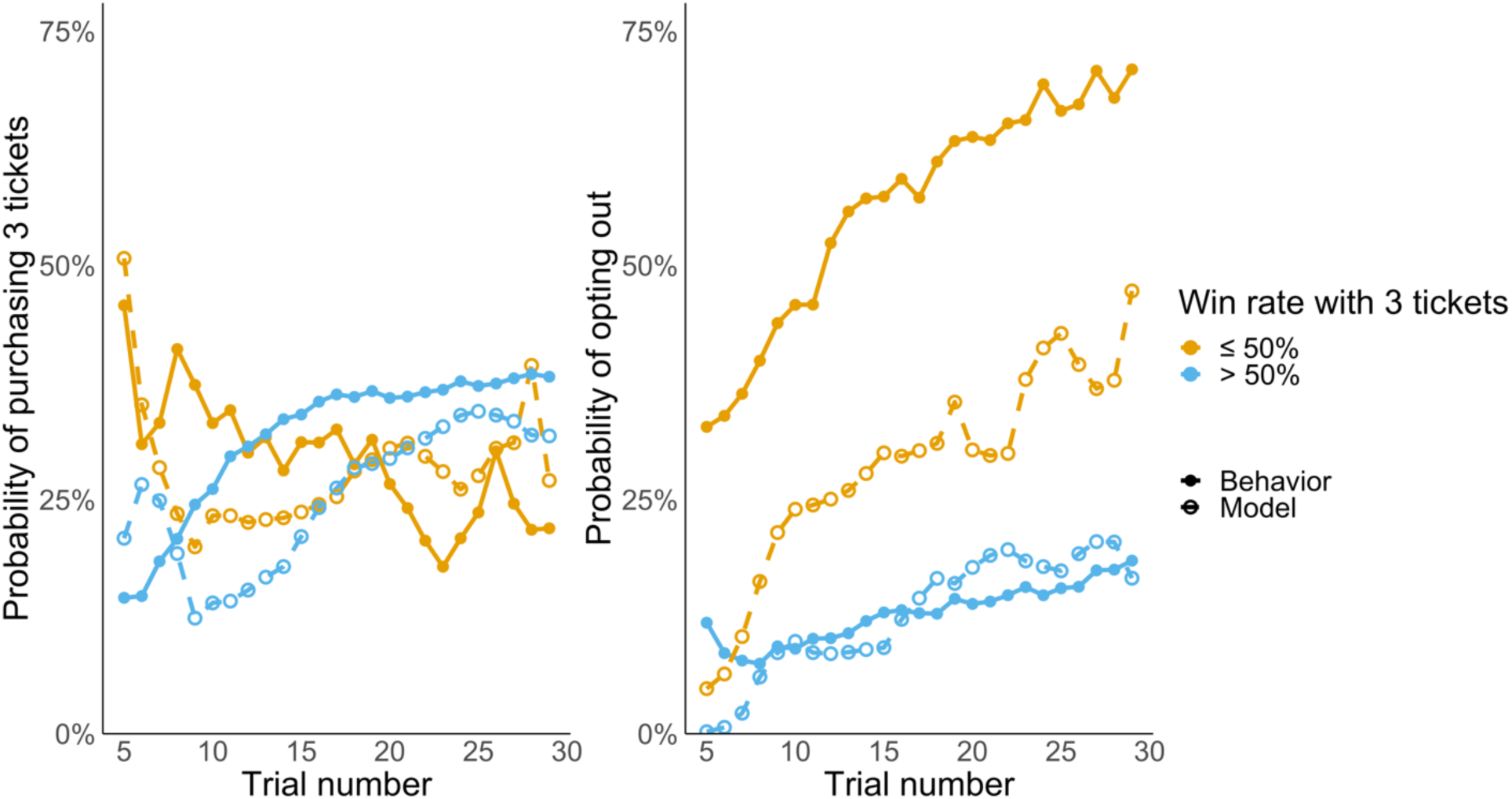
Behavioral trajectories under high and low controllability conditions. Left: Probability of investing maximum resources (3 tickets) across trials. **Right:** Probability of opting out. Data points are divided by experienced win rates with 3 tickets (orange: ≤50%, blue: >50%). Solid lines with filled circles represent participant behavior; dashed lines with empty circles show computational model predictions. Participants experiencing low controllability initially persist with maximum investment before gradually shifting to opting out as negative evidence accumulates.

**Supplementary Figure 3.**
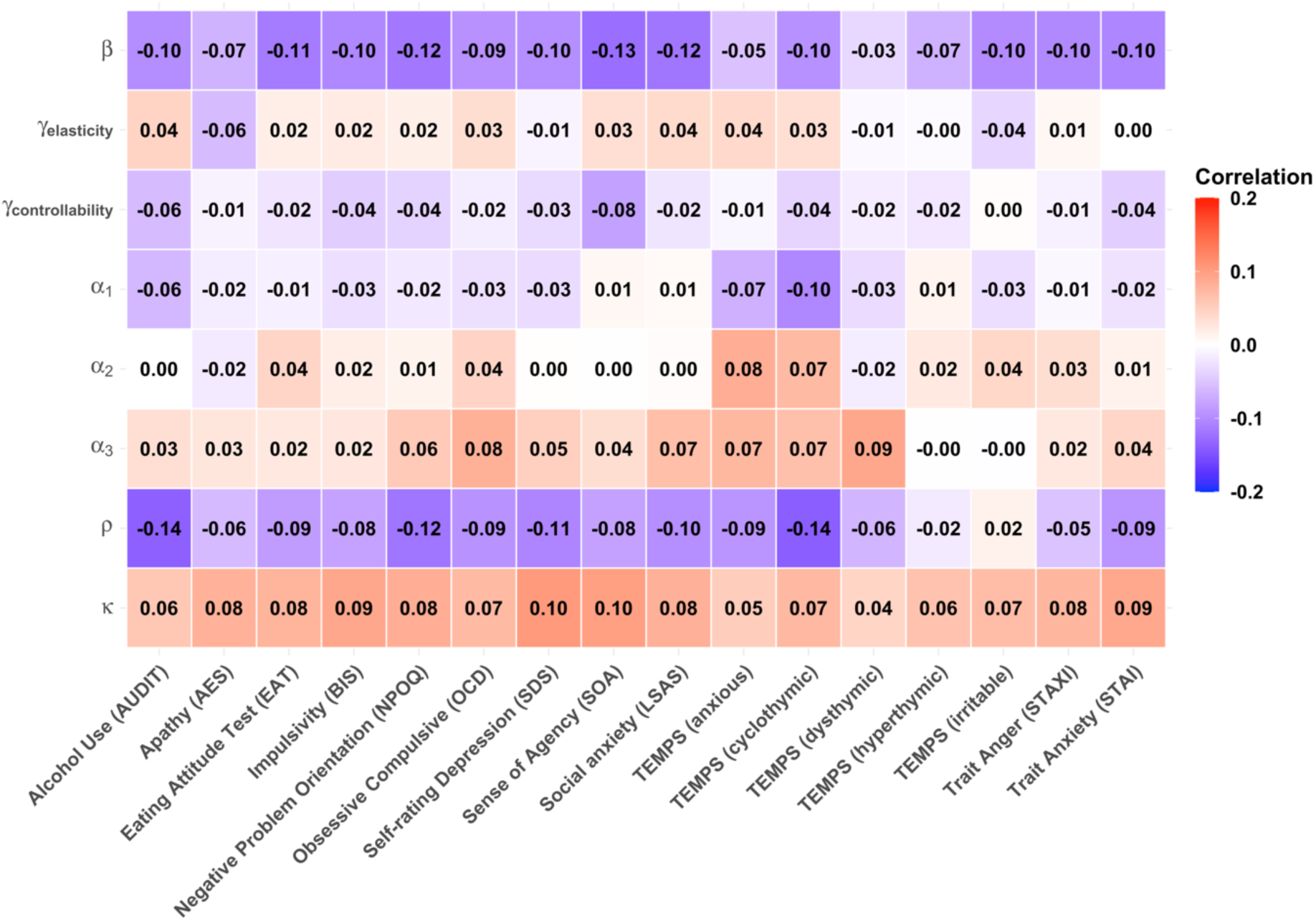
Correlation coefficients between model parameter and questionnaire scores. Pearson correlation of computational model parameters (rows) with psychological questionnaire scores (columns). Color intensity indicates correlation strength, with blue representing negative correlations and red representing positive correlations. No correlation reached statistical significance when correcting for multiple comparisons (FDR).

**Supplementary Figure 4.**
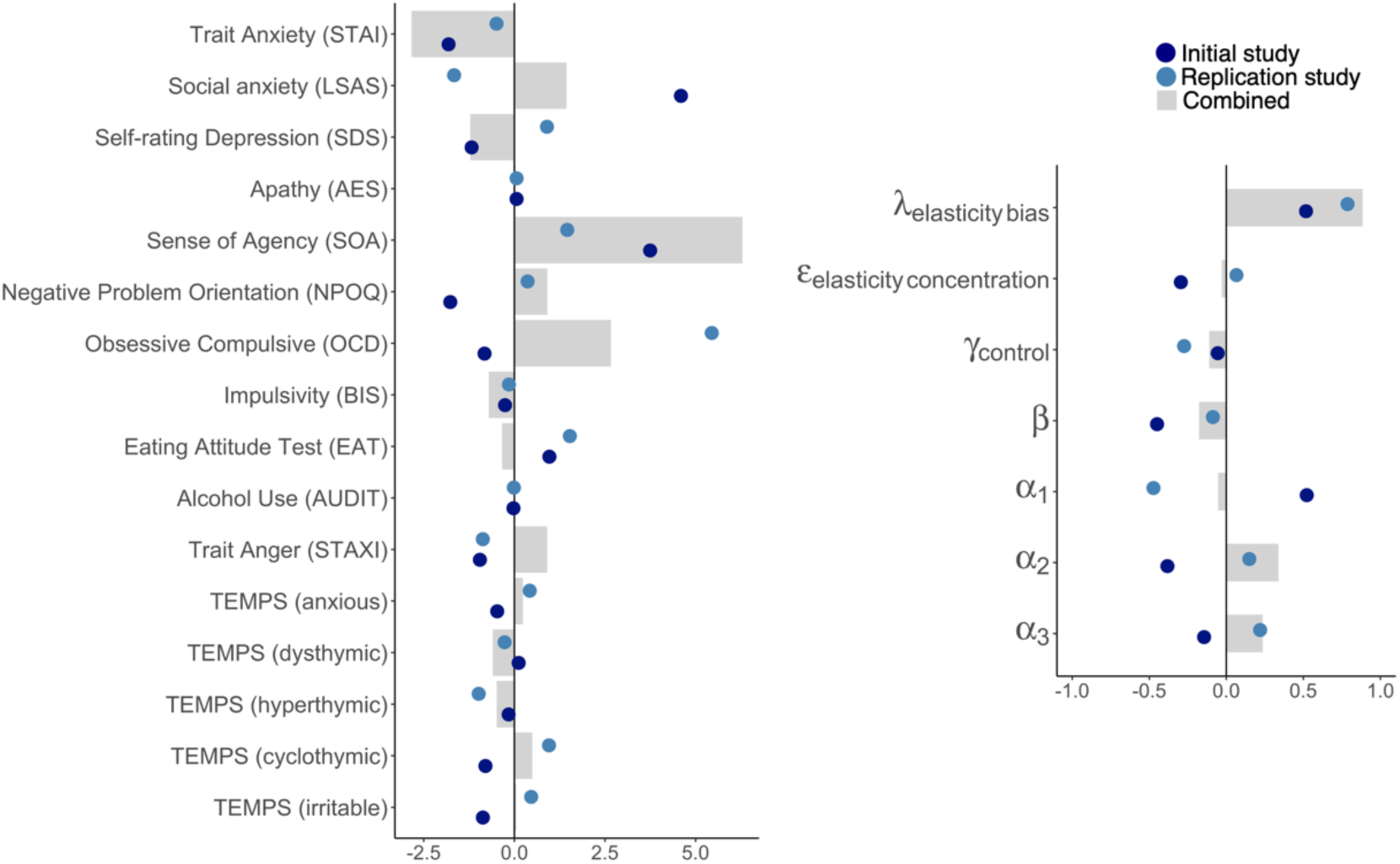
Model parameters and psychopathology. Loadings from a Canonical Correlation Analysis (CCA) between model parameters and self-report psychopathology measures while separating concentration from bias parameters in the elasticity prior. Self-report measures were scored such that higher values reflect higher levels of psychopathology. Loadings are shown for the Initial (dark blue), Replication (light blue), and combined (gray bars) datasets. The magnitude of the loading reflects the degree to which each measure contributes to the correlation between parameters and self-reports

